# Exploit Spatially Resolved Transcriptomic Data to Infer Cellular Features from Pathology Imaging Data

**DOI:** 10.1101/2024.08.05.606654

**Authors:** Zhining Sui, Ziyi Li, Wei Sun

## Abstract

Digital pathology is a rapidly advancing field where deep learning methods can be employed to extract meaningful imaging features. However, the efficacy of training deep learning models is often hindered by the scarcity of annotated pathology images, particularly images with detailed annotations for small image patches or tiles. To overcome this challenge, we propose an innovative approach that leverages paired spatially resolved transcriptomic data to annotate pathology images. We demonstrate the feasibility of this approach and introduce a novel transfer-learning neural network model, STpath (Spatial Transcriptomics and pathology images), designed to predict cell type proportions or classify tumor microenvironments. Our findings reveal that the features from pre-trained deep learning models are associated with cell type identities in pathology image patches. Evaluating STpath using three distinct breast cancer datasets, we observe its promising performance despite the limited training data. STpath excels in samples with variable cell type proportions and high-resolution pathology images. As the influx of spatially resolved transcriptomic data continues, we anticipate ongoing updates to STpath, evolving it into an invaluable AI tool for assisting pathologists in various diagnostic tasks.

## Introduction

Manual pathological examination of histology images remains the gold standard for evaluating and diagnosing many diseases, such as cancer [1], neurodegenerative diseases [2], and heart failure [3]. Hematoxylin and eosin (H&E)-stained whole slide images (WSIs) are the most common type of histology images. They are gigapixel images that often contain over a billion pixels each, making manual inspection highly labor-intensive. As routine screening and examination become prevalent, the volume of WSI data is growing faster than the number of proficient pathologists available. An artificial intelligence (AI)-assisted WSI analysis system, primarily driven by deep learning methods, offers a promising solution to this shortage of pathology resources [4]. Such an AI system can have an even greater impact in low-income regions or developing countries, where pathology expertise is scarce. Additionally, it can enhance interobserver reproducibility, which is crucial for improving diagnostic consistency [5].

Building such a deep learning system requires a large amount of high-quality training data. Manual examination of WSIs follows a hierarchical approach: starting with the whole image, identifying the area of interest, and zooming into small patches for detailed analysis. Therefore, the training data needs to be annotated at the level of these small patches (e.g., 224 × 224 pixel squares). However, most available H&E-stained WSIs are only labeled at the whole image level, and manually annotating these WSIs at the patch level is prohibitively laborious.

Recent advancements in spatially resolved transcriptomics (SRT) offer unprecedented opportunities to study gene expression in cells within their native tissue microenvironment. In contrast to WSIs, SRT provides much richer information that can be used to estimate cell types or cell states. Typically, a WSI is collected alongside SRT data from the same tissue sample, creating the potential to annotate the WSI using the annotations derived from SRT data. Extracting annotations from SRT data is an active research area and many computational methods have been developed to call cell types of individual cells or estimate cell type composition at each spot on the WSI [6, 7, 8].

In this paper, we present a transfer learning approach to develop a convolutional neural network (CNN) capable of annotating WSIs. Starting with a pre-trained CNN, we retrain it using training data that includes paired WSIs and SRT data. The trained model can then be applied to make predictions on testing data where only WSI data are available. Despite the limited amount of training data, we demonstrate the feasibility and good generalizability of our method. We have developed a pipeline named STpath (Spatially resolved Transcriptomics and pathology imaging data) and applied it to two types of tasks: predicting cell type proportions and classifying tumor microenvironments. We also provide guidance on best practices, such as selecting hyperparameters for CNNs, and options for working with images with relatively lower resolutions.

## Methods

### SRT data

We applied the proposed STpath pipeline to four SRT datasets, one from a melanoma brain metastasis and three from breast cancer samples. The SRT data for brain metastasis (patient 16) was generated by Sudmeier et al. [9] using the 10x Visium platform, from a 10 µm thick H&E-stained flash-frozen section of the tumor parenchyma. The slide was imaged with a Lionheart Microscope (Biotek) at 10x magnification.

The first BRCA dataset was publicly shared by 10x Genomics. It includes two BRCA tissue samples generated using the 10x Visium platform. The full-resolution WSIs provided by 10x Genomics were scanned using a 20X magnification microscope objective. The second BRCA dataset was generated by Wu et al. [10] using the 10x Visium solution, consisting of six human breast cancer samples. The third BRCA dataset, generated by He et al. [11], consists of samples from 23 patients. For each patient, three microscope images of H&E-stained tissue slides were taken. Corresponding SRT data were collected using spatial transcriptomics, which has lower resolution than 10x Visium.

### Image processing

The SRT data provide the position and diameter of each gene expression spot. Using this information, we generated *standard* square patches, with each patch covering one spot. The size of each patch, in terms of the number of pixels, varied across datasets: 50 × 50 pixels for the brain metastasis dataset [9], 90× 90 pixels for the four CID-samples of Wu et al. [10], 152 *× 1*52 pixels for the 23 samples of He et al. [11], and around 200 × 200 pixels for all other samples. Such variation was due to the difference in imaging capture resolution and magnification. To facilitate unified processing, each image was rescaled to 224 × 224 pixels for input into the CNN.

The resolutions of the extracted patches for each sample tissue used in this study are shown in Supplementary Table 1. Additionally, we generated *large* patches for the datasets obtained using the 10x Visium solution. These *large* patches encompassed eight spots within each patch (Supplementary Figure 1). Since the Visium spots are hexagonally spaced, the arrangement of these eight spots followed a specific pattern, forming an hourglass shape. Specifically, there were three spots in the first and third rows of the large patch, while the second row contained two spots.

Some of the gene expression spots do not overlap with the tissue sample. When extracting patches from WSIs, we only utilized spots within the tissue that contained gene expression data. Specifically, only patches containing at least 50% tissue were selected for further analysis. To distinguish tissue from the background, we adopted a multi-step approach similar to the one used by Barker et al. [12]. First, we converted the patch images to grayscale, merging the three color channels into a single channel, with each pixel assigned an integer value from 0 to 255. Since the slide backgrounds were illuminated with white light, the pixel values in the grayscale image’s background were mostly close to or equal to 255. Next, we enhanced the contrast of the grayscale image using adaptive histogram equalization [13], ensuring better differentiation of tissue from the background. The complement of the contrast-enhanced grayscale image was then obtained at an 8-bit depth, inverting the pixel values so that the background values were close to or equal to 0. Finally, we performed hysteresis thresholding, an automatic edge detection technique, to detect tissue boundaries. Briefly, hysteresis thresholding detects edges using the following rules: edges with intensities above the high threshold are identified as true edges; edges with intensities below the low threshold are classified as non-edges; and edges with intensities between the high and low thresholds are identified as edges only if they are connected to a previously identified true edge; otherwise, they are discarded. Other contrast enhancement techniques, such as contrast stretching, and thresholding techniques, such as Otsu’s, could also be implemented. Supplementary Figure 2 provides examples of tissue detection, using patches from different sources.

### Stain normalization

Stain normalization is performed to address the variability in staining intensity and color that can arise from differences in staining protocols, reagents, and imaging conditions. These variations can introduce biases and affect the accuracy of image analysis. We employed two methods for stain normalization: the Macenko method [14] and the Vahadane method [15]. These algorithms transfer the color style of the source image to that of a carefully selected target image while preserving other information in the processed image. Supplementary Figure 3 provides two examples of stain normalization, one from sample 1142243F and another from sample CID4465. In our analysis, neither stain normalization method improved STpath’s performance in prediction or classification. Therefore, we report the results generated using the original images unless otherwise noted. It is possible that stain normalization may be more beneficial for larger and more heterogeneous cohorts.

### Cell type deconvolution for the regression task

We employed a cell type deconvolution algorithm called CARD (Conditional AutoRegressive model-based Deconvolution) to estimate the composition of cell types in spatial transcriptomic data [16]. CARD takes advantage of the spatial information available in the data and performs reference-based deconvolution. The relevant cell types of a tissue need to be determined for each tissue type separately, and then scRNA-seq data of these cell types are needed as input for CARD. CARD provides estimates of the proportions of relevant cell types within each spot.

We did not run CARD on the SRT data retrieved from Sudmeier et al. [9] due to the lack of scRNA-seq reference data with annotations for relevant cell types. We used a breast cancer scRNA-seq dataset generated by Wu et al. [10] as the reference for cell type deconvolution of all BRCA datasets. This dataset consists of scRNA-seq (Chromium, 10x Genomics) data from 26 primary tumors representing three major clinical subtypes of breast cancer: 11 ER+, 5 HER2+, and 10 TNBC tumors. The scRNA-seq data identified nine major cell types and 29 minor cell types. We ran CARD on the three SRT datasets. The deconvolution results are consistent with annotations for breast cancer samples from the 10x Genomics dataset and Wu et al. [10] dataset, but not for the He et al. [11] data, possibly due to lower resolution of the He et al. data.

### Class annotation for the classification task

For the brain metastasis data [9] and the BRCA data from Wu et al. [10], the clusters were generated and annotated by the authors (Supplementary Table 3-4). For the BRCA dataset from He et al. [11], we clustered the gene expression spots using Seurat V5 [17] (Supplementary Table 6). We log-normalized the expression data and integrated data from different samples using anchor-based CCA (Canonical Correlation Analysis) integration.

### Neural Network

ResNets are a popular choice for supervised transfer learning due to their superior performance and efficiency compared to other CNN models [18]. In this study, we used neural networks built upon a ResNet-50 architecture pre-trained on the ImageNet dataset, though other pre-trained models can also be used in the STpath pipeline.

ResNet-50, as the name implies, consists of 50 layers. We excluded the final output layer, originally containing 1,000 nodes for ImageNet classification tasks. To preserve the learned weights in the base model, we froze these layers to prevent further weight updates. To adapt the base model to our specific tasks, we introduced new trainable layers on top of it. These additional layers converted the extracted features from the base model into predictions aligned with our regression or classification tasks. The size of the final output layer, a Fully Connected (FC) layer, was determined based on the nature of each task. For regression tasks, the number of nodes in the final output layer matched the number of cell types we aimed to predict. For classification tasks, the number of nodes corresponded to the number of distinct classes within the dataset. The output layer used the softmax activation function for all tasks, allowing our model to generate predictions compatible with the output expectations of each task.

Between the final output layer and the base model, we introduced an additional FC layer with the ReLU activation function. This hidden FC layer increased the number of trainable parameters, thereby enhancing the model’s capacity. To mitigate the risk of overfitting, we incorporated a dropout layer immediately after the intermediate FC layer. We fine-tuned the dropout proportion and the number of nodes in the hidden FC layer to optimize performance.

### Training and Evaluation

While we trained different networks for different datasets and tasks, we consistently divided the available patches into three subsets: 70% for training, 15% for validation, and 15% for testing. To facilitate comparison across neural networks with different hyperparameters, the division of the training, validation, and testing datasets remained consistent for each dataset.

To train the models, we employed different optimizers, including Adam, SGD, and RMSprop, aiming to minimize categorical entropy loss for classification tasks and mean squared error loss for regression tasks. We experimented with varying learning rates and batch sizes to enhance the model’s learning. The optimal hyperparameters were identified based on performance improvements on the validation dataset.

We saved the model’s weights after each epoch if there was an improvement in loss on the validation dataset. The maximum number of training epochs was set to be 1,000 for all CNN models. Additionally, we implemented early stopping based on the validation loss, with a patience of 30 epochs. If the validation loss did not decrease within 30 epochs, we terminated the training process.

To evaluate the model’s performance on the testing dataset, we used metrics appropriate for the task. For classification tasks, we utilized AUC, while for regression tasks, we measured the mean absolute error and mean squared error.

## Results

### An overview of STpath

*An overview of STpath. (A) An illustration to split a WSI into individual patches. The left panel shows the WSI of a breast cancer FFPE sample downloaded from the 10x genomics website. The other panels illustrate the process of splitting the WSI into patches. (B) Generate patch annotations or features such as tissue microenvironment or cell type proportions. (C) Given image patches* **X** *and features associated with each patch* **y**, *use transfer learning to build a CNN to learn features from image patches*.

The transfer-learning-based STpath pipeline consists of three steps. First, a WSI is segmented into small patches (Fig. 1(A)). By default, each patch contains one spot of SRT data and is referred to as a *standard* patch. Additionally, *large* patches are created, each encompassing eight spots. We use patches that contain at least 50% tissue, selected through a three-step process similar to the one described by Barker et al. [12]. To mitigate color and intensity biases in stained images, we apply stain normalization.

**Fig. 1.**
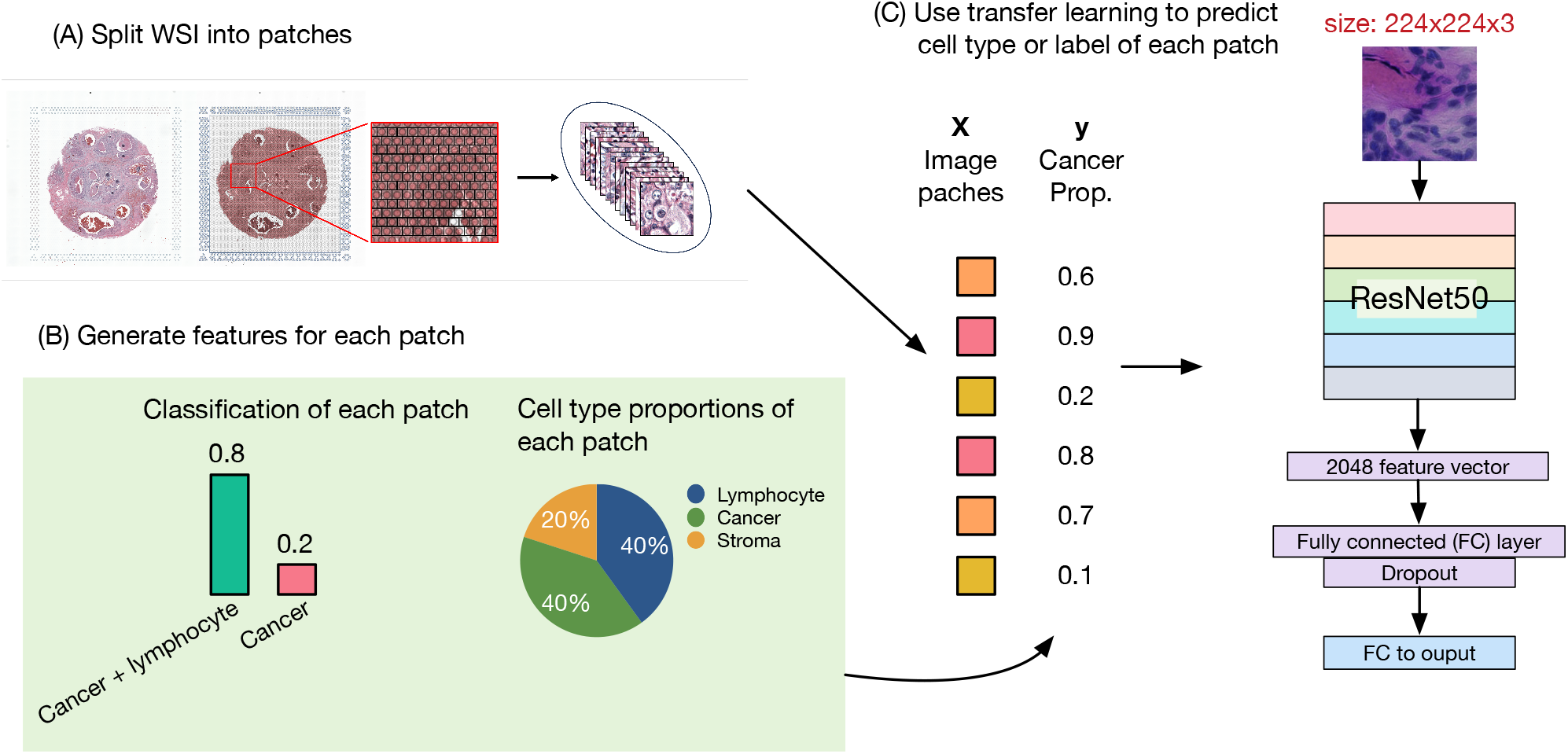
An overview of STpath. (A) An illustration to split a WSI into individual patches. The left panel shows the WSI of a breast cancer FFPE sample downloaded from the 10x genomics website. The other panels illustrate the process of splitting the WSI into patches. (B) Generate patch annotations or features such as tissue microenvironment or cell type proportions. (C) Given image patches **X** and features associated with each patch **y**, use transfer learning to build a CNN to learn features from image patches.

After creating and pre-processing the image patches, the second step involves generating features for each patch, which the CNN will predict (Fig. 1(B)). These features can include proportions of cell types inferred by deconvolution methods from SRT data [19, 20, 21, 22, 16] or class labels derived from SRT data [23, 24] or pathologists. STpath supports both regression and classification tasks. For regression, the goal is to quantitatively predict the cell type proportions within each patch. For classification, the objective is to categorize each patch into distinct tissue microenvironments.

In the final step, we use transfer learning to train a CNN to make predictions on the features of interest for image patches (Fig. 1(C)). In this work, the pre-trained baseline model is ResNet50 [18], though it can be replaced by any other pre-trained CNNs. ResNet50 utilizes residual blocks that include multiple shortcut connections, adding the input back to the output and enabling the network to learn residual features (Supplementary Figure 4). We modify ResNet50 by appending a hidden layer of 256 neurons to the ResNet50 output, followed by a dropout layer and a fully connected layer to generate the output (Fig. 1(C)). Further details of our method and pipeline are provided in the Methods section.

### A proof-of-concept example

We began with an example involving the classification of two spatial regions in a melanoma brain metastasis sample [9] with an H&E image and paired 10x Visium SRT data. The authors identified 7 clusters using the SRT data: tumor (cluster 4), tumors adjacent to inflammation (cluster 5), inflammation (cluster 1), and inflammation with blood vessels (cluster 7) (Fig. 2(A)).

**Fig. 2.**
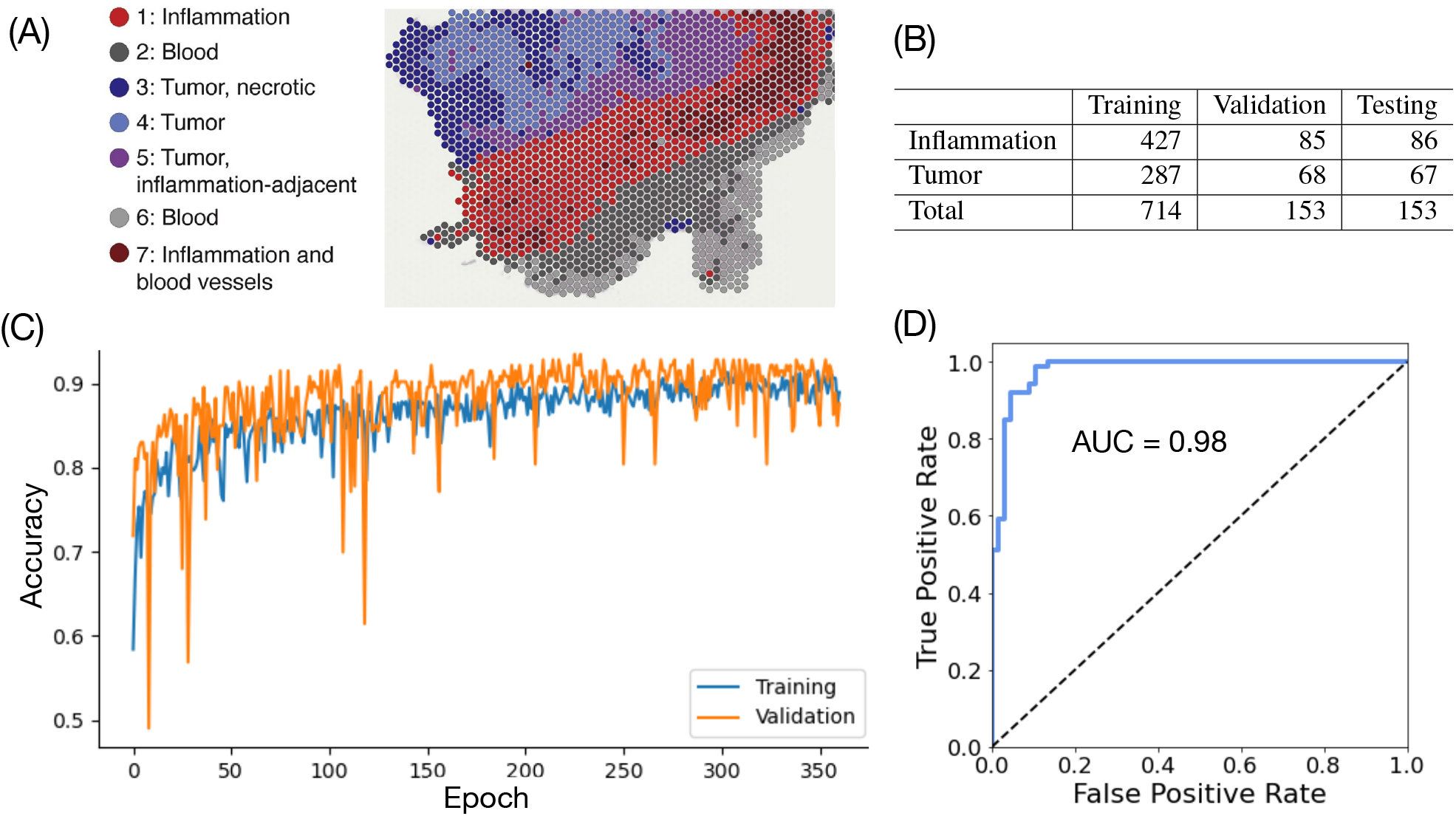
(A) Cluster assignments of spots from an SRT slide of a melanoma brain metastasis sample. This panel is a modified version of Figure 6b of Sudmeier et al. [9], which is an open-access article under the CC BY license. (B) Sample size for training, validation, and testing. (C) Classification accuracy in training and validation data along training epochs. (D) Classification ROC curve in testing data.

Three other clusters included two clusters of blood and one cluster of necrotic tumors. Our focus was on classifying tumors (clusters 4 and 5) versus inflammation (clusters 1 and 7). Although this classification task was relatively simple due to the homogeneity of each region, it also posed a challenge because of the limited amount of training data. The dataset comprised approximately 1,000 spots: around 700 for training, 150 for validation, and 150 for testing (Fig. 2(B)).

Using this example data, STpath achieved high classification accuracy. The training of STpath concluded after roughly 360 epochs. The validation accuracy was approximately 0.9 with no sign of overfitting (Fig. 2(C)). The AUC (area under the receiver operating characteristic curve) for the testing data was around 0.95 (Fig. 2(D)). We also explored two stain normalization methods: the Macenko method [14] and the Vahadane method [15]. The accuracy of the classification before and after either stain normalization was similar.

This proof-of-concept example demonstrated that STpath, trained with only *∼*700 spots, could achieve accurate classification.

### Applications to breast cancer datasets

In the remaining part of the paper, we explore paired WSI and SRT data from breast cancer (BRCA) samples, which are more representative tumor tissues with mixed cell types. The BRCA samples came from three sources: two samples from 10x Genomics, six samples generated by Wu et al. [10], and 23 samples generated by He et al. [11].

The first 10x sample was derived from an FFPE human breast tissue obtained from BioIVT Asterand Human Tissue Specimens and was annotated as Ductal Carcinoma In Situ (DCIS) and Invasive Carcinoma (Fig. 3(A)). The second 10x sample was derived from a fresh frozen Invasive Ductal Carcinoma breast tissue, also obtained from BioIVT (Figure 3(B)).

**Fig. 3.**
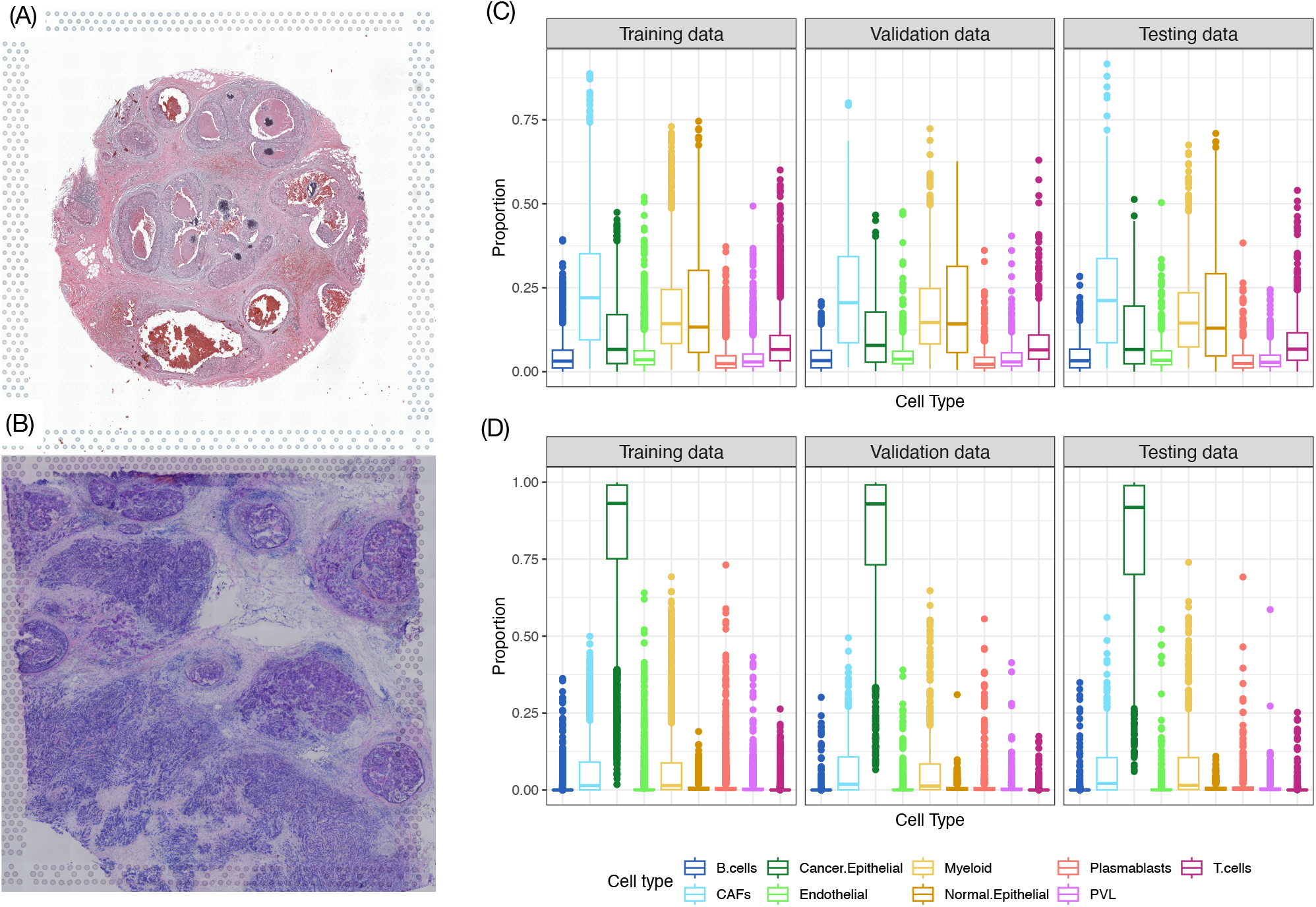
(A) The WSI of the 10x FFPE sample. (B) The WSI of the 10x fresh frozen sample. (C) Cell type proportion estimates for all the spots of the FFPE sample. (D) Cell type proportion estimates for all the spots of the fresh frozen sample. CAF: cancer-associated fibroblast. PVL: perivascular-like stroma cells.

The SRT dataset from Wu et al. [10] consisted of six human breast samples, including two samples classified as ER+ (CID4535 and CID4290), two classified as triple-negative breast cancers (TNBC) (CID44971 and CID4465), and two additional TNBC samples processed in an independent laboratory (1142243F and 1160920F). The authors generously provided full-resolution WSIs of all six tissues upon request. The first four samples were at 20x magnification, while the last two were at 40x magnification. The cell type compositions varied significantly across these samples (Table 1, Supplementary Figure 5(A)).

**Table 1.** Number of patches in each tissue section with each of the nine major cell types having the highest proportion.

| Cell Type | 10x Genomics |  | Wu et al. [2021] |  |  |  |  |  |
| --- | --- | --- | --- | --- | --- | --- | --- | --- |
|  | FFPE | Fresh Frozen | 1142243F | 1160920F | CID4290 | CID4465 | CID44971 | CID4535 |
| B cells | 13 | 18 | 547 | 878 | 531 | 46 | 694 | 14 |
| CAFs | 980 | 49 | 545 | 0 | 324 | 1008 | 78 | 51 |
| Cancer Epithelial | 127 | 4516 | 3144 | 61 | 1556 | 65 | 46 | 946 |
| Endothelial | 49 | 43 | 0 | 0 | 2 | 0 | 2 | 1 |
| Myeloid | 431 | 225 | 22 | 1 | 1 | 21 | 19 | 43 |
| Normal Epithelial | 736 | 0 | 19 | 273 | 0 | 12 | 137 | 0 |
| Plasmablasts | 8 | 33 | 406 | 3681 | 2 | 56 | 176 | 54 |
| PVL | 17 | 13 | 0 | 1 | 16 | 1 | 10 | 4 |
| T cells | 157 | 1 | 98 | 0 | 0 | 2 | 0 | 14 |

The dataset from He et al. [11] included 23 human breast samples. These samples were classified into five categories: five HER2-luminal, five HER2-non-luminal, four Luminal A, five Luminal B, and four TNBC samples. For each sample, three replicated WSIs were obtained, together with the corresponding SRT data.

The SRT data from the two 10x samples and the Wu et al. [10] samples were generated by the 10x Visum platform, where each slide captured an area of 6.5 × 6.5 mm and approximately 5000 spots. Each spot was 55 µm in diameter with a 100 µm center to center distance and each spot covered 1-10 cells. The SRT data from He et al. [11] had relatively lower resolution where each spot had a diameter of 100 µm with a 200 µm center-to-center distance.

To estimate the cell type composition of each spot, we employed the computational method CARD [16], which showed good performance in independent evaluations [8]. We used a breast cancer scRNA-seq dataset [10] as the reference for cell type deconvolution. The dataset consisted of scRNA-seq data from 26 primary tumors representing three major clinical subtypes of breast cancer: 11 ER+ cancers, 5 HER2+ cancers, and 10 TNBCs. CARD revealed nine major cell types. Cell type proportions varied across samples. For example, in the 10x FFPE sample, multiple cell types were present, with cancer-associated fibroblasts (CAFs) being the most abundant one (Fig. 3(C)). Conversely, the 10x fresh frozen sample predominantly consisted of tumor cells (Fig. 3(D)). Several cell types had low proportions across patches, making them harder to study. Therefore, in the following analysis, we also considered four collapsed cell types that were defined as follows: Invasive Cancer (Cancer Epithelial), Stroma (Endothelial, PVLs, or CAFs), Lymphocyte (T cells, B cells, or Plasmablasts), and Others (Normal Epithelial or Myeloid) (Supplementary Figure 5(B)).

In addition to cell type proportions, we also generated class labels for classification tasks. We adopted the class labels reported by Wu et al. [10] and generated class labels for the SRT data from He et al. [11] by clustering all spots from all samples.

### Features learned by ResNet

ResNet was originally trained on ImageNet, which consists of a wide variety of images significantly different from H&E-stained WSIs. We first posed a question: Could the 2,048 features extracted by the pre-trained ResNet capture cellular information from WSIs? We generated four patch groups for each tissue sample from the two 10x samples and the Wu et al. samples. Specifically, Groups 1 and 2 had comparable proportions of a specific cell type, as did Groups 3 and 4. However, the proportions of this specific cell type differed between Groups 1 and 2 versus Groups 3 and 4. Next, we compared the values of each ResNet50-extracted feature between any two groups using the rank-sum test. We assessed the difference between the two groups by calculating the proportion of features with p-values smaller than 0.05. If there was no difference between the two groups, this proportion should have been approximately 0.05, and higher proportions indicated that more features differed than expected.

There were six pairwise comparisons for the four groups, with two being within-category comparisons (Group 1 vs. 2, and Group 3 vs. 4) and four being between-category comparisons. Due to the compositional nature of cell type proportions (i.e., the proportions of the four cell types added up to 1), it was not possible to adjust the proportion of one cell type while leaving the proportions of other cell types unchanged. Thus, we simply sampled the patches with high or low proportions of one cell type and did not put constraints on other cell types (Supplementary Figures 6-10). The results were challenging to interpret because a ResNet feature might capture the signal from any cell type. Nevertheless, it was clear that many ResNet features were different for between-category comparisons, and around 5% of ResNet features were different for within-category comparisons, which was expected to happen by chance (Fig. 4).

**Fig. 4.**
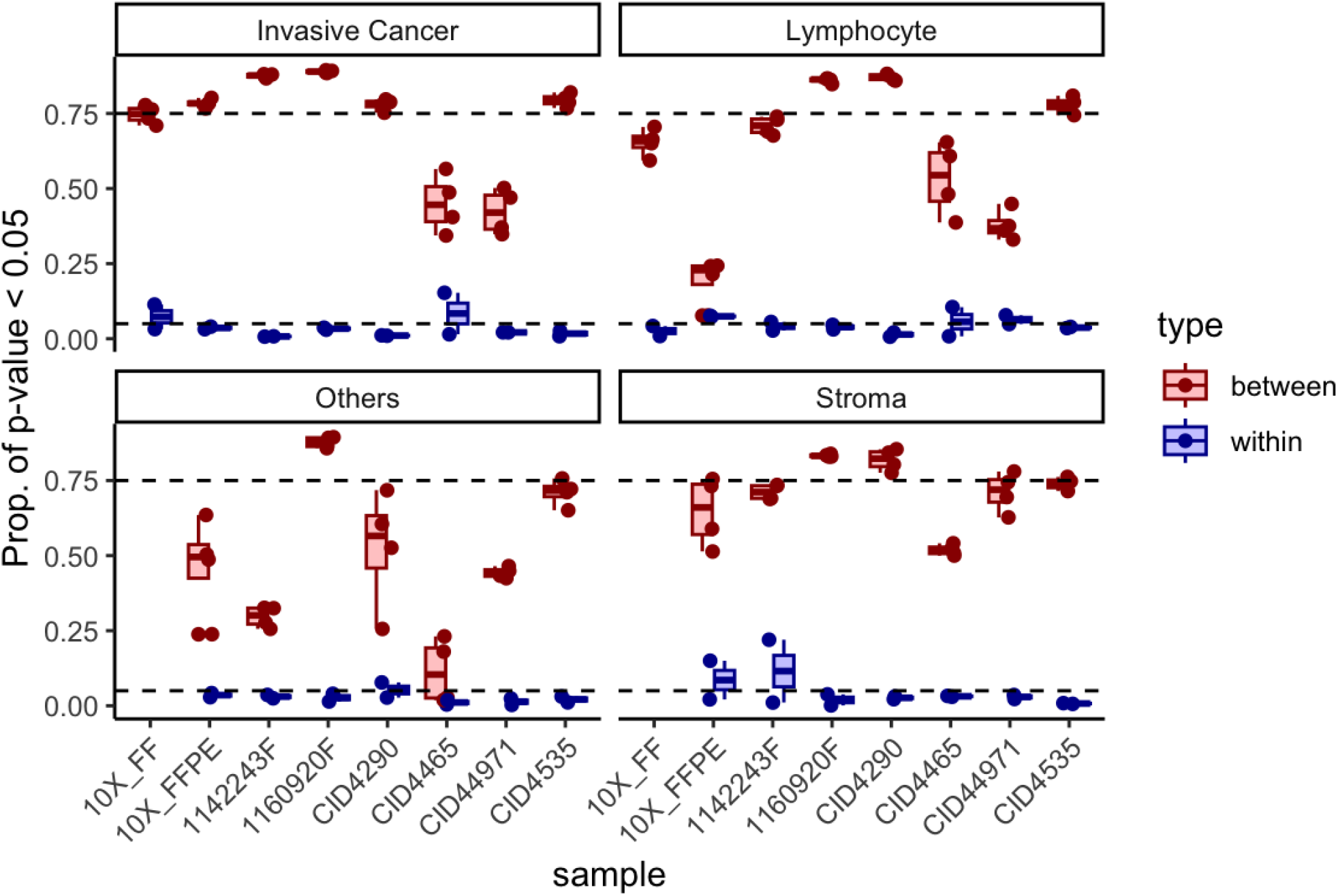
The proportion of rank-sum test p-values less than 0.05, when comparing the 2048 ResNet features between groups defined by cell type proportions and samples. The two horizontal lines indicated 0.05 and 0.75, respectively.

If a cell type had relatively lower proportions in the spots of one slide, sampling spots with higher or lower proportions of this cell type did not change the proportions of other cell types much. This created the opportunity to evaluate the identified features associated with a particular cell type. For example, the results from CID4465 and CID44971 were more informative in identifying features associated with invasive cancer proportions (Supplementary Figure 7), CID44971 was an informative sample for lymphocytes (Supplementary Figure 8), CID4465 was an informative sample when comparing proportions of other cell types (epithelia and myeloid) or stroma (Supplementary Figures 9-10).

The above results demonstrated that the pre-trained ResNet could recover features of WSIs that were associated with cell type identity. Next, we evaluated the modified ResNet with additional layers trained with paired WSI and SRT data.

### Prediction of cell type proportion

Cell identity is one of the most important molecular features when reading a H&E WSI. We sought to predict cell type proportions for each patch of a WSI. We constructed a *standard* patch around a spot of paired SRT data. Each dataset was split into training, validation, and testing sets. The evaluation was performed on the testing data using the hyperparameters selected by the validation data.

For the two 10x Genomics samples, we first trained STpath using two samples together. The predicted cell type proportions in the testing data showed strong correlations with the observed proportions estimated by CARD (Fig. 5(A)). The cell type compositions are very different between the two samples. The 10x fresh frozen sample predominantly consisted of cancer cells, while the 10x FFPE sample contained a mixture of several cell types. This disparity could introduce a batch effect, potentially biasing the results of the CNN. We also trained STpath separately for each sample. Training the model separately for each sample yielded higher prediction accuracy in the testing data for some cell types, although the overall performance remained similar (Supplementary Figure 11).

When applying the same procedure to Wu et al. data [10], we observed clear batch effects impacting prediction accuracy. The predictions for the two 11-samples were positively correlated with estimates from the SRT data. In contrast, STpath performed poorly on the four CID-samples; the predicted values exhibited little variation and were clustered around their mean values (Fig. 5(B)). We conjectured that this issue was due to the low resolution of the CID-samples, where a *standard* patch was approximately 90 × 90 pixels. In contrast, a *standard* patch of the 11-samples was around 200 × 200 pixels. To address this discrepancy, we constructed *large* patches covering eight spots in the SRT data (See Methods section for details). Despite the reduction in sample size for training due to larger patches, the prediction accuracy improved for the 11-samples and the combined 11-and CID- samples (Fig. 5(C)). Although the performance is still poor for the CID- samples, the improved *R*^2^’s in all samples suggest that larger patches help align the two sets of samples. However, suboptimal performance on the CID- samples suggests that further resolution-related adjustments may be required.

**Fig. 5.**
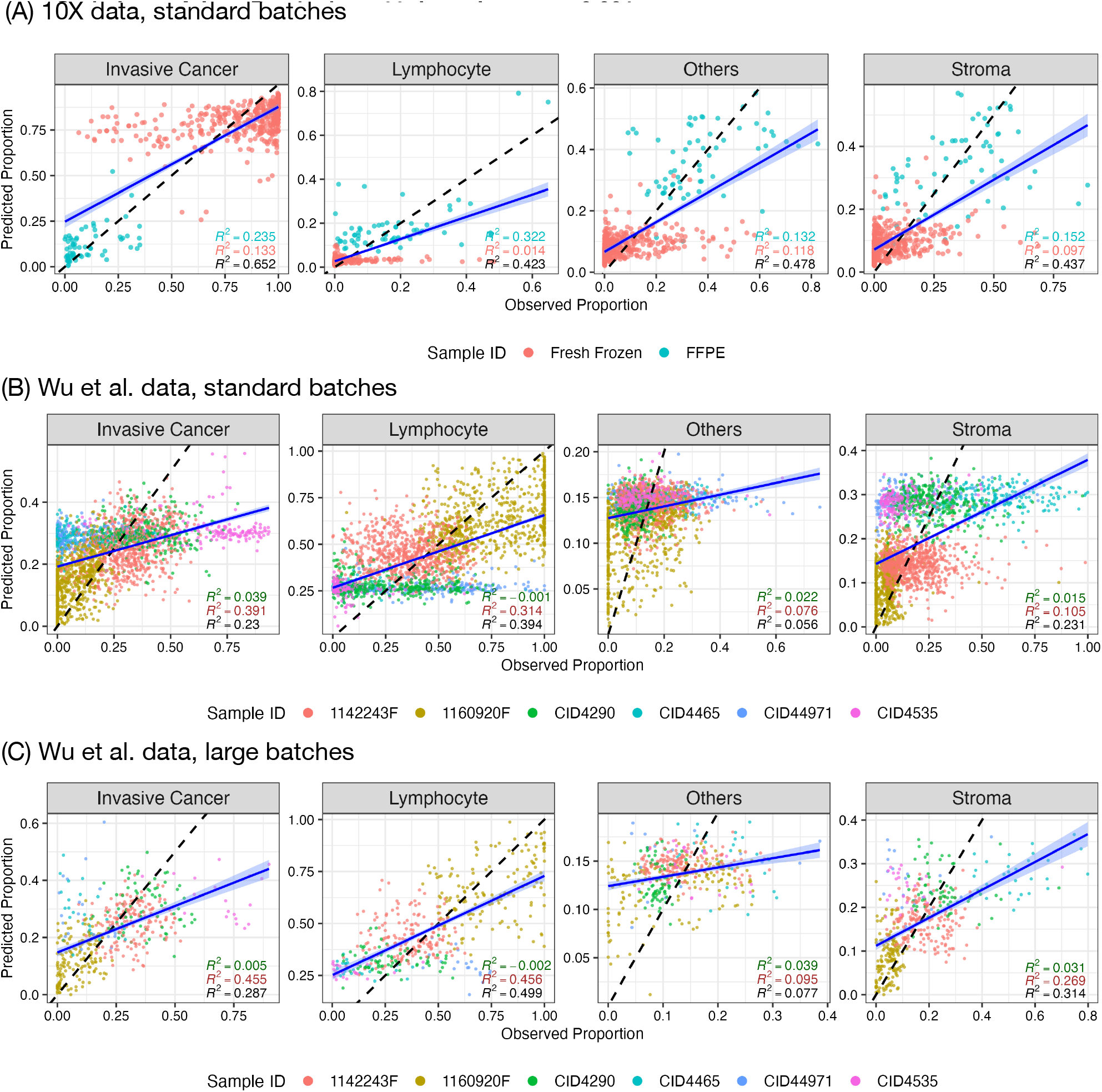
Comparison of observed cell type proportions estimated by CARD versus predicted cell type proportions by STpath using testing data and the hyperparameters selected by validation data. (A) Comparison for two 10x samples using standard patches (i.e., each patch represents a spot) with the hyperparameters: Batch size (batch) = 32, learning rate (LR) = 0.001, dropout proportion (dropout) = 0.0, and size of dense layer (size) = 512. Three *R*^2^s for Fresh frozen (red), FFPE (blue), and combined data (black) were shown for each cell type. (B) Comparison for six samples from Wu et al. [10] using using standard patches with the hyperparameters: batch = 32, LR = 0.0001, dropout = 0.0, and size = 512. Three *R*^2^s for CID-(dark green), 11-(dark red), and combined data (black) were shown for each cell type. (C) Comparison for six samples from Wu et al. [10] using large patches such that one patch covers eight spots. The hyperparameters are as follows: batch = 128, LR = 0.0001, dropout = 0.2, and size = 512.

We also trained STpath models to predict the proportion of nine cell types. The prediction results for nine cell types had similar patterns as those for four cell types (Supplementary Figures 12-13). We could estimate the proportions of four cell types by collapsing the prediction results of the 9-cell-type model. Interestingly, the predictions obtained through this post-collapsing approach were very similar to those derived from directly predicting the proportions of the four cell types (Supplementary Figure 14).

### Classification of tumor microenvironment

In the proof-of-concept example, we considered a classification problem and STpath achieved high accuracy when classifying Tumor vs. Inflammation cell regions. In this section, we extended our exploration of classification tasks to more challenging scenarios where each class was defined by a specific tumor microenvironment. We utilized annotations from Wu et al. [10], which classified each spot of the SRT data into one of the following 11 classes: adipose tissue, invasive cancer, invasive cancer + lymphocytes, invasive cancer + stroma, invasive cancer + stroma + lymphocytes, lymphocytes, necrosis, normal + stroma + lymphocytes, normal glands + lymphocytes, stroma, and stroma + adipose tissue. Patches annotated as NA, Artefact, or Uncertain were excluded from the analysis. Additionally, for each sample, labels that were assigned to fewer than 50 patches were also excluded to ensure sufficient data representation for each class. We computed the AUC for each class, as well as the aggregated AUCs across all classes.

The performance of STpath varied across samples and classes within each sample (Fig. 6(A), Supplementary Figure 15). Overall, the performance was better in the two 11-samples compared to the four CID-samples. This is consistent with our findings for predicting cell type proportions, suggesting that the relatively low resolution of WSIs from CID-samples affects classification accuracy. For the two 11-samples, the most challenging classification was for stroma, with AUCs around 0.74. The tumor tissues in these two samples were included in the class “Invasive cancer + stroma + lymphocytes”. This class was classified with high accuracy, achieving AUCs of 0.832 and 0.872 for samples 1142243F and 1160920F, respectively.

We also performed classifications on the He et al. dataset [11], with annotations generated by clustering via Seurat v5 [17]. We detected three clusters, denoted by clusters 0, 1, and 2, and they had an increasing proportion of spots annotated as tumors based on the tumor/non-tumor annotations provided by He et al [11] (Fig. 6(B)). The fact that all clusters have a considerable amount of tumor may reflect the preference to select tumor regions for this SRT data. As expected, it was easier to distinguish the spots in cluster 0 or cluster 2 from other spots because they had relatively low or high tumor content, while it was harder to isolate cluster 1 because it had intermediate tumor content (Fig. 6(C)).

**Fig. 6.**
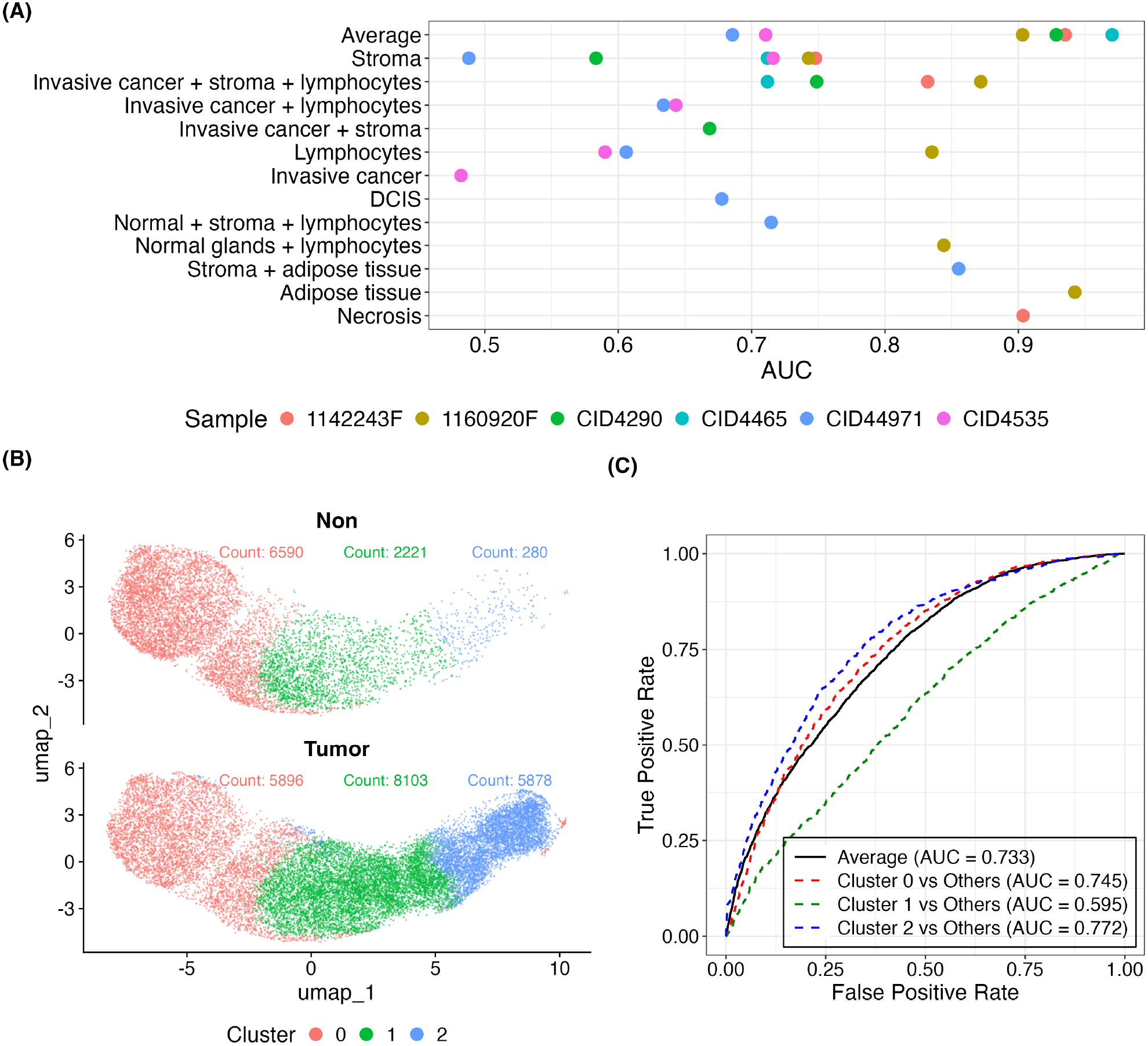
(A) Micro-averaged and class-specific AUCs for the classification task in the testing data of six samples from Wu et al. [10]. The optimal neural network configuration was selected using validation data, with the following hyperparameters: optimizer = Adam, LR = 0.0001, dropout = 0.0, size = 512, and batch size = 32 for CID4465, and 16 for all other samples. (B) UMAP visualization of 28,968 gene expression spots analyzed by RNA expression from SRT data and integrated across 23 breast tumors from He et al. [11] using the CCA method, stratified by He et al.’s tumor/non-tumor annotations. UMAP dimensional reduction was performed using 30 principal components in the Seurat v5 package. Clusters were created with a resolution of 0.1. The number of spots in each cluster is displayed, colored by cluster. (C) ROC curve for the classification task in the testing data of 23 samples from He et al. [11]. The optimal neural network configuration was selected using validation data, with the following hyperparameters: batch = 256, optimizer = Adam, LR = 0.001, dropout = 0.0, and size = 256.

## Discussion and conclusion

Deep learning methods hold great promise for extracting valuable information from whole slide images (WSIs) to predict clinical outcomes [25, 26] and molecular traits [27]. However, a significant challenge in training these methods is obtaining patch-level annotations.

A popular approach that does not require patch-level annotations is multi-instance learning [25, 26]. It treats a WSI as a bag of patches and classifies the bag if at least one patch meets the criteria for a specific class. This assumption works well for certain tasks, such as distinguishing between tumor and non-tumor samples, where the presence of a single tumor patch suffices to classify the WSI as a tumor sample. However, this assumption may not hold for many other tasks, such as studying the tumor-immune microenvironment.

The STpath framework proposed in this paper differs fundamentally from multi-instance learning. STpath leverages spatially resolved transcriptomic data to provide batch-level labels for model training. Once trained, an STpath model can be applied to WSIs to translate a large (e.g., billions of pixels) and shallow (e.g., two color channels for H&E-stained images) WSI into a smaller (e.g., thousands of patches) and deeper (e.g., composition of nine cell types) image. This can greatly simplify downstream analysis, facilitating the assessment of associations between WSIs and clinical outcomes. Additionally, this approach enhances interpretability by providing biologically meaningful annotations for each patch.

While it seems intuitive to use paired WSI and SRT to train a neural network, key questions remain. These include determining the number of samples needed, identifying appropriate neural network structures, and deciding which molecular features to derive from the SRT data. Our work addresses these fundamental questions. We discovered that numerous features derived from ImageNet-pre-trained neural networks were associated with cell type labels of image patches. To the best of our knowledge, our group is the first to report these results, which strongly support the use of transfer learning in studying WSIs. Additionally, this explains our observation that a neural network with good generalizability can be trained with just a few thousand patches.

We have explored a wide range of hyperparameters for the neural networks, including the optimizer, batch sizes, learning rate, the size of the hidden layer that connects the pre-trained ResNet features with the output layer, and the dropout rate for the dropout layer following the hidden layer. Our findings indicate Adam optimizers with a learning rate of 0.001 or 0.0001 tend to outperform other optimizers. The results are generally not sensitive to other hyperparameters, which can be selected by validation data. We recommend choosing a batch size of 32, 64, or 128, a hidden layer with 256 or 512 hidden nodes, and a dropout rate of 0, 0.2, or 0.5.

Using three breast cancer datasets, we aimed to predict two types of molecular features for breast cancer: cell type proportions and tumor microenvironments. Both tasks could be achieved with relatively high accuracy. The cases with poorer performance were samples predominantly of one cell type or those with low-resolution WSIs. Previous research has focused on predicting the expression of individual genes from H&E-stained images [11]. An interesting future direction would be to explore how gene expression prediction is mediated by the prediction of cell types.

A major challenge of using paired WSI and SRT data to train STpath is the noise in molecular features derived from SRT due to estimation uncertainty, which could be due to the uncertainty of extracting molecular features by a computational method or the resolution of the SRT data. For example, using higher resolution SRT data [10], we can obtain more refined clusters than with lower resolution SRT data [11]. There are a few directions to mitigate this estimation uncertainty. One direction is to employ more advanced neural network training methods that can account for noise in the training data [28, 29, 30]. Another direction is to use SRT data with cellular resolution, for example, the SRT data from the recent CosMx platform [31].

In this work, all the prediction and classification tasks were conducted for each patch separately. However, leveraging information from nearby patches could potentially improve STpath’s performance. This is challenging because the pattern of spatial dependence can vary greatly across tissue types. Ideally, this problem should be formulated as an image segmentation problem. By dividing a WSI into different spatial regions, information can be borrowed within those regions. Therefore, we envision an iterative framework that alternates between prediction/classification and image segmentation tasks.

## Key Points

- Spatially resolved transcriptomic (SRT) data can be used to generate both continuous features (e.g., cell type proportions) and discrete features (e.g., classes of tumor microenvironment) for paired H&E stained images.
- Transfer-learning is an effective solution to train deep learning method to interpret H&E stained images. When applying pre-trained deep learning models to H&E stained images, the derived features are strongly associated with cell type labels.
- Higher resolution of SRT data and H&E stained images leads to better performance.

## Supporting information

Supplementary results

## Data availability

The codes for STpath are available at https://github.com/Sun-lab/STpath with detailed instructions on its usage. The main data supporting the results of this study are available within the paper and its Supplementary Information. The 10x Visium SRT data for brain metastases is publicly available through the Gene Expression Omnibus under accession number GSE179572. The SRT data for the 10x FFPE and the 10x fresh frozen BRCA samples can be downloaded from https://www.10xgenomics.com/resources/datasets/human-breast-cancer-ductal-carcinoma-in-situ-invasive-carcinoma-ffpe-1-standard-1-3-0 and https://www.10xgenomics.com/resources/datasets/human-breast-cancer-visium-fresh-frozen-whole-transcriptome-1-standard, respectively. All 10x Visium SRT data in Wu et al. are available from the Zenodo data repository (https://doi.org/10.5281/zenodo.4739739). All SRT data in He et al. are available at https://data.mendeley.com/datasets/29ntw7sh4r/5. The scRNA-seq reference used for cell type deconvolution is publicly available through the Gene Expression Omnibus under accession number GSE176078.

