## Supplementary results for "Exploit Spatially Resolved Transcriptomic Data to Infer Cellular Features from Pathology Imaging Data"

### Supplementary Information for "Exploit Spatial Transcriptomic Data to Infer Cellular Features from Pathology Imaging Data"

Zhining Sui, Ziyi Li, Wei Sun

### 1 Supplementary Information for Methods

#### 1.1 SRT data introduction

In general, there are two types of Spatially Resolved Transcriptomics (SRT) data: Spatial Transcriptomics (ST) and 10x Visium. ST combines high-resolution tissue imaging with high-throughput transcriptome sequencing. A key feature of the ST method is its ability to simultaneously capture histological imaging and gene expression data. Histological imaging uses standard staining techniques to detail the tissue's structure, while gene expression profiling processes and sequences spatially barcoded cDNA to determine gene activity. These gene expression datasets are then aligned with the images to ensure accurate visualization. The primary advantage of ST is its unbiased ability to capture mRNA abundance and spatial location, enabling the redefinition of morphological features based on molecular characteristics. The initial version of ST microarrays consisted of approximately 1,000 spots with a diameter of 100 µm and a center-to-center distance of 200 µm, over an area of 6.2 mm by 6.6 mm.

In 2018, 10x Genomics enhanced and commercialized ST technology as Visium Spatial Gene Expression, increasing sensitivity and throughput. 10x Genomics Visium is a next-generation molecular profiling solution for spatial gene expression, enabling measurement of the entire transcriptome in a spatially resolved manner by mapping gene expression onto high-resolution microscope images of intact fresh frozen or FFPE tissue sections. The Visium Gene Expression slide features either two tissue capture areas measuring 11 mm $\times$ 11 mm or four areas measuring 6.5 mm $\times$ 6.5 mm. A standard 6.5 mm $\times$ 6.5 mm capture area covers approximately 5,000 barcoded gene expression spots, while an XL 11 mm $\times$ 11 mm capture area covers approximately 14,000 barcoded spots. Compared to ST, the spot size is reduced from 100 µm to 55 µm in diameter, and capture area density is enhanced with hexagonal packing, having a distance of 100 µm between the centers of adjacent spots. A WSI is typically magnified 20 or 40 times. At 40$\times$ magnification, one square micrometer is represented by 4 $\times$ 4 pixels, resulting in an image of 6.5 mm $\times$ 6.5 mm containing approximately 0.68 billion pixels.

| | Image capture magnification | Dimensions of WSI (pixel× pixel) | Dimensions of patches (pixel× pixel) |
|---|---|---|---|
| 10X Genomics FFPE Breast Tissue | 20x objective | 27452 × 25233 | Standard: 208 × 208 |
| 10X Genomics Fresh Frozen Breast Tissue | 20x objective | 41572 × 27755 | Standard: 194 × 194 |
| Sample CID4290 Processed by Wu et al. [1] | 20x objective | 9907×9404 | Standard: 90 × 90<br>Large: 290 × 290 |
| Sample CID4465 Processed by Wu et al. [1] | 20x objective | 9648 × 9232 | Standard: 90 × 90<br>Large: 290 × 290 |
| Sample CID44971 Processed by Wu et al. [1] | 20x objective | 9648 × 9232 | Standard: 90 × 90<br>Large: 290 × 290 |
| Sample CID4535 Processed by Wu et al. [1] | 20x objective | 9648 × 9232 | Standard: 90 × 90<br>Large: 290 × 290 |
| Sample 1142243F Processed in An Independent Laboratory | 40x objective | 41572 × 27755 | Standard: 224 × 224<br>Large: 720 × 720 |
| Sample 1160920F Processed in An Independent Laboratory | 40x objective | 41572 × 27755 | Standard: 226 × 226<br>Large: 724 × 24 |
| Melanoma Brain Metastasis Processed by Sudemeier et al. | 10x objective | 15796 × 14355 | Standard: 50 × 50 |
| 23 Samples Processed by He et al. | 20x objective | ∼ 9300× 9900 | Standard: 160 × 160 |

Supplementary Table 1: Summary of resolution and size in WSIs and patches. The samples were scanned using various commercial WSI systems from different manufacturers. It should be noted that different WSI scanning models may have distinct image capture resolutions (microns/pixel) at the same image capture magnification, resulting in varying dimensions of WSIs under identical image capture magnification.

#### 1.2 Image preprocessing

*Standard* patches were created such that each patch contained only one SRT spot. The radius of each patch was set to be slightly larger than the radius of each spot (1.1 times larger) to ensure there was no overlap between different patches. However, other folds can be used. For the 10x Visium platform, the radius of a spot is 27.5 µm, and the distance between the centers of two spots should be 100 µm. Therefore, the radius of the patches can be at most 1.8 times larger than the spots to avoid overlap between patches. For data obtained by the ST platform, the radius of a spot is 50 µm, and the distance between the centers of two spots should be 200 µm. Thus, the radius of the patches can be at most two times larger than the spots to avoid overlap between patches. Additionally, large patches were created for 10x Visium data, as shown in Supplementary Figure 1.

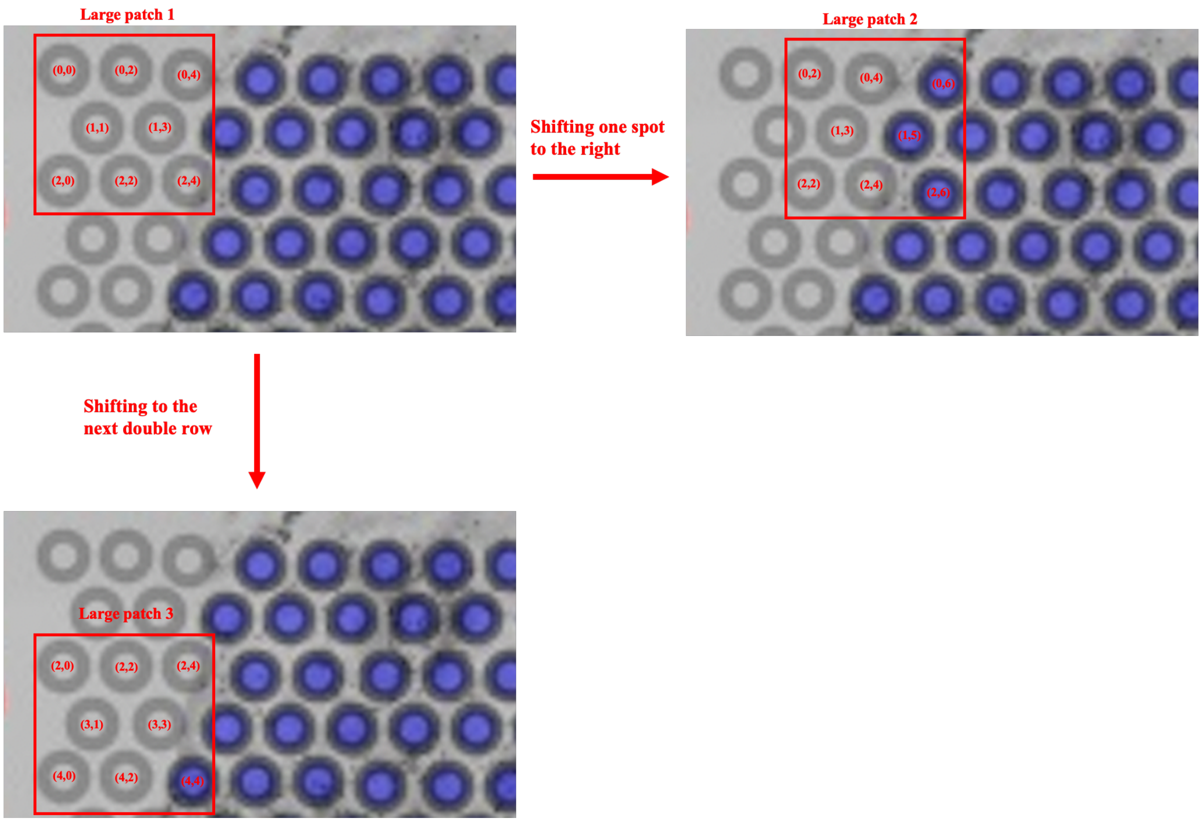

Supplementary Figure 1: Process of generating large patches.

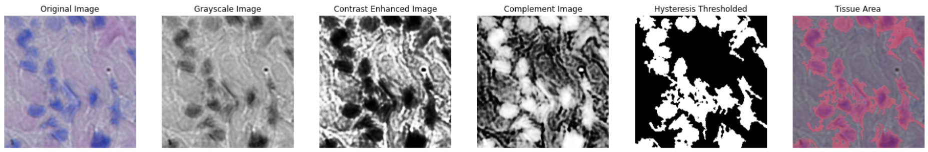

(A) A patch from 10x fresh frozen sample.

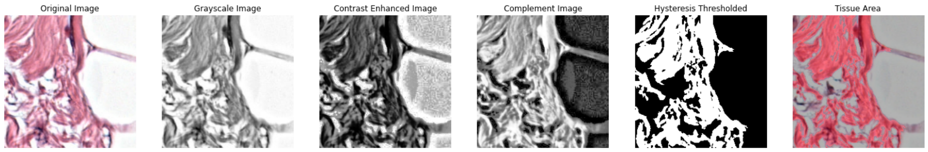

(B) A patch from 10x FFPE sample.

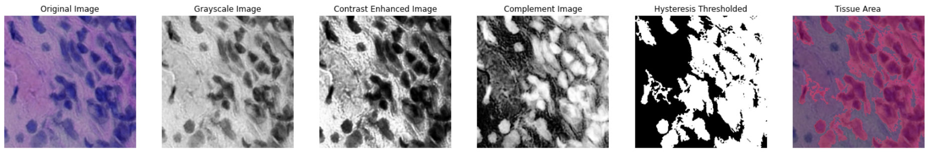

(C) A patch from Sample 1142243F from Wu et al. [1].

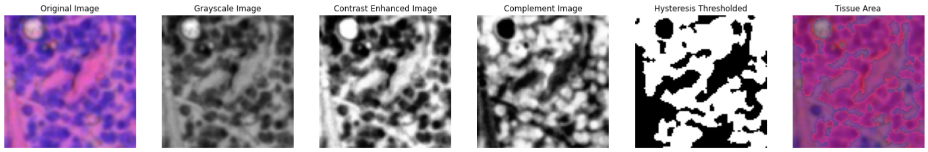

(D) A patch from Sample CID4465 from Wu et al. [1].

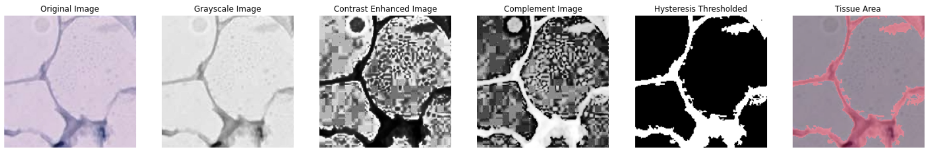

(E) A patch from Sample BC23287 from He et al. [2].

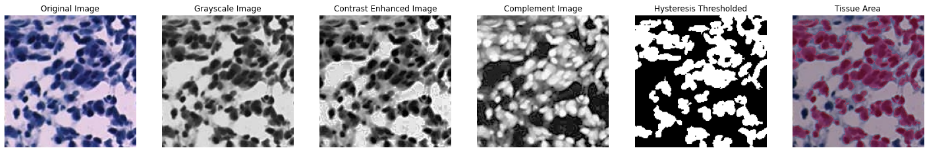

(F) A patch from Sample BC23377 from He et al. [2].

Supplementary Figure 2: Examples of quality check for patches using tissue detected.

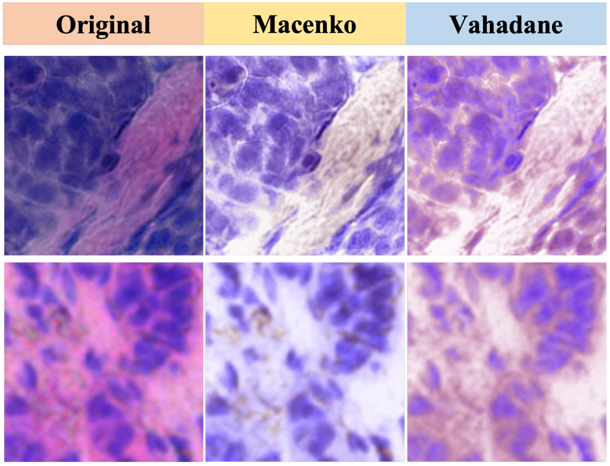

Supplementary Figure 3: Examples of stain normalization.

| | Number of patches | | |
|---|---|---|---|
| | Training | Validation | Testing |
| 1142243F | 581 | 144 | 169 |
| 1160920F | 637 | 172 | 154 |
| CID4290 | 292 | 75 | 72 |
| CID4465 | 178 | 47 | 27 |
| CID44971 | 203 | 39 | 20 |
| CID4535 | 150 | 33 | 23 |

Supplementary Table 2: Number of large patches from each of the six samples from Wu et al. [1] in the training, validation, and testing datasets.

#### 1.3 CNN structure

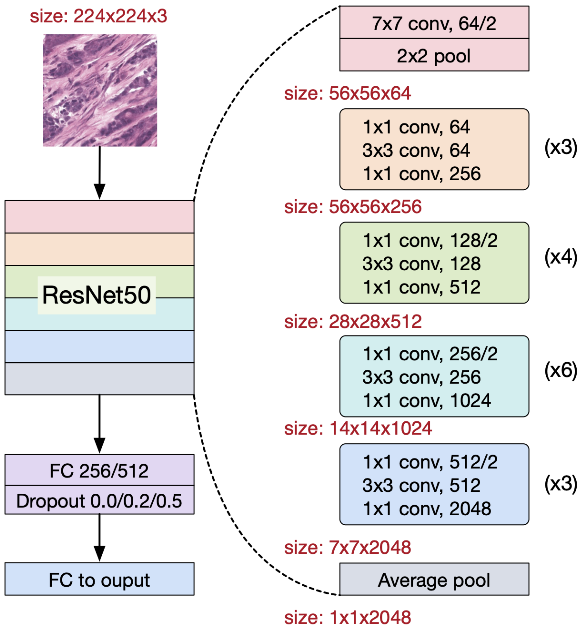

Supplementary Figure 4: CNN architecture overview.

#### 1.4 Annotations for classification

| Cluster | Number of Spots |
|---|---:|
| 1: Inflammation | 451 |
| 2: Blood | 289 |
| 3: Tumor, necrotic | 231 |
| 4: Tumor | 218 |
| 5: Tumor, inflammation-adjacent | 204 |
| 6: Blood | 176 |
| 7: Inflammation and blood vessels | 147 |

Supplementary Table 3: Number of SRT spots from Sudmeier et al. [3] within each cluster defined by the authors.

| | 1142243F | 1160920F | CID4290 | CID4465 | CID44971 | CID4535 | Total |
|---|---|---|---|---|---|---|---|
| Adipose tissue | 0 | 83 | 0 | 0 | 0 | 8 | 91 |
| Artefact | 119 | 48 | 7 | 4 | 1 | 23 | 202 |
| Cancer trapped in lymphocyte aggregation | 0 | 9 | 0 | 0 | 0 | 0 | 9 |
| Ductal carcinoma in situ (DCIS) | 0 | 12 | 0 | 0 | 273 | 0 | 285 |
| Invasive cancer | 0 | 0 | 0 | 0 | 0 | 418 | 418 |
| Invasive cancer + adipose tissue + lymphocytes | 0 | 0 | 0 | 0 | 0 | 3 | 3 |
| Invasive cancer + lymphocytes | 0 | 0 | 0 | 0 | 317 | 361 | 678 |
| Invasive cancer + stroma | 0 | 0 | 2082 | 0 | 0 | 0 | 2082 |
| Invasive cancer + stroma + lymphocytes | 3627 | 3146 | 215 | 1131 | 0 | 0 | 8119 |
| Lymphocytes | 15 | 186 | 0 | 0 | 81 | 69 | 351 |
| Necrosis | 568 | 0 | 0 | 0 | 0 | 0 | 568 |
| Normal + stroma + lymphocytes | 0 | 0 | 0 | 0 | 240 | 0 | 240 |
| Normal duct | 0 | 0 | 0 | 3 | 0 | 0 | 3 |
| Normal glands + lymphocytes | 0 | 278 | 0 | 0 | 0 | 0 | 278 |
| Stroma | 445 | 1132 | 122 | 73 | 134 | 169 | 2075 |
| Stroma + adipose tissue | 0 | 0 | 0 | 0 | 114 | 0 | 114 |
| Tertiary lymphoid structures (TLS) | 10 | 0 | 0 | 0 | 0 | 0 | 10 |
| Uncertain | 0 | 0 | 0 | 0 | 0 | 73 | 73 |
| NA | 0 | 1 | 6 | 0 | 2 | 3 | 12 |

Supplementary Table 4: Number of SRT spots in each patient from Wu et al. [1] with each pathology annotation.

| | Training | Validation | Testing |
|---|---|---|---|
| **1142243F** | | | |
| Invasive cancer + stroma + lymphocytes | 2531 | 537 | 559 |
| Necrosis | 396 | 97 | 75 |
| Stroma | 321 | 62 | 62 |
| **1160920F** | | | |
| Adipose tissue | 57 | 15 | 11 |
| Invasive cancer + stroma + lymphocytes | 2218 | 463 | 465 |
| Lymphocytes | 123 | 32 | 31 |
| Normal glands + lymphocytes | 195 | 44 | 39 |
| Stroma | 785 | 170 | 177 |
| **CID4290** | | | |
| Invasive cancer + stroma | 1454 | 319 | 309 |
| Invasive cancer + stroma + lymphocytes | 158 | 23 | 34 |
| Stroma | 81 | 21 | 20 |
| **CID4465** | | | |
| Invasive cancer + stroma + lymphocytes | 789 | 170 | 172 |
| Stroma | 54 | 10 | 9 |
| **CID44971** | | | |
| DCIS | 194 | 42 | 37 |
| Invasive cancer + lymphocytes | 214 | 49 | 54 |
| Lymphocytes | 60 | 12 | 9 |
| Normal + stroma + lymphocytes | 162 | 39 | 39 |
| Stroma | 101 | 18 | 15 |
| Stroma + adipose tissue | 80 | 14 | 20 |
| **CID4535** | | | |
| Invasive cancer | 296 | 67 | 55 |
| Invasive cancer + lymphocytes | 249 | 49 | 63 |
| Lymphocytes | 45 | 11 | 13 |
| Stroma | 122 | 25 | 22 |

Supplementary Table 5: Patch counts for each of the six samples from Wu et al. [1], categorized by the annotations involved in the classification task. DCIS: Ductal carcinoma in situ

| | He et al. | | Seurat v5 | | |
|---|---|---|---|---|---|
| | Non | Tumor | Cluster 0 | Cluster 1 | Cluster 2 |
| BC23209 | 43 | 857 | 297 | 321 | 282 |
| BC23268 | 1078 | 259 | 642 | 468 | 227 |
| BC23269 | 200 | 1093 | 646 | 287 | 360 |
| BC23270 | 0 | 815 | 353 | 342 | 120 |
| BC23272 | 515 | 1126 | 874 | 441 | 326 |
| BC23277 | 413 | 1148 | 538 | 239 | 784 |
| BC23287 | 405 | 235 | 215 | 270 | 155 |
| BC23288 | 127 | 1285 | 640 | 717 | 55 |
| BC23377 | 457 | 1421 | 968 | 29 | 881 |
| BC23450 | 86 | 808 | 332 | 494 | 68 |
| BC23506 | 84 | 1192 | 273 | 790 | 213 |
| BC23508 | 588 | 833 | 735 | 287 | 399 |
| BC23567 | 878 | 818 | 886 | 699 | 111 |
| BC23803 | 51 | 834 | 350 | 100 | 435 |
| BC23810 | 248 | 1839 | 656 | 900 | 531 |
| BC23895 | 66 | 1361 | 170 | 1050 | 207 |
| BC23901 | 279 | 259 | 271 | 182 | 85 |
| BC23903 | 1003 | 260 | 730 | 400 | 133 |
| BC23944 | 200 | 1006 | 672 | 173 | 361 |
| BC24044 | 1305 | 206 | 907 | 511 | 93 |
| BC24105 | 0 | 954 | 356 | 474 | 124 |
| BC24220 | 107 | 1170 | 408 | 688 | 181 |
| BC24223 | 958 | 98 | 567 | 462 | 27 |

Supplementary Table 6: Number of SRT spots from He et al. [2] within each cluster defined by the authors or generated by Seurat v5 [4].

The macro-averaged AUCs were calculated by determining the AUC for each class individually and then averaging them, treating all classes equally. In contrast, the micro-averaged AUCs were obtained by aggregating the contributions of all samples, giving equal weight to each sample. In our multi-class classification setup, the micro-average was preferred and reported due to class imbalance.

### 2 Additional results

#### 2.1 Estimates of cell type proportions using SRT data

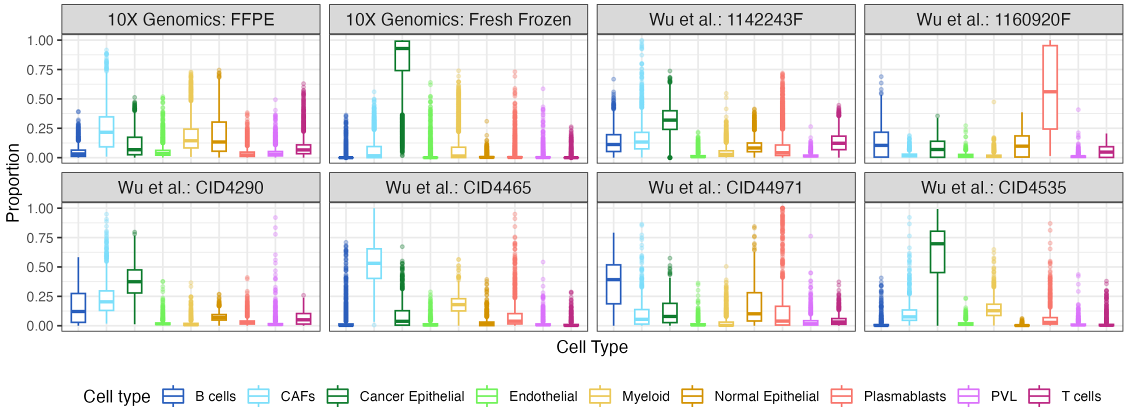

(A) Nine major cell types.

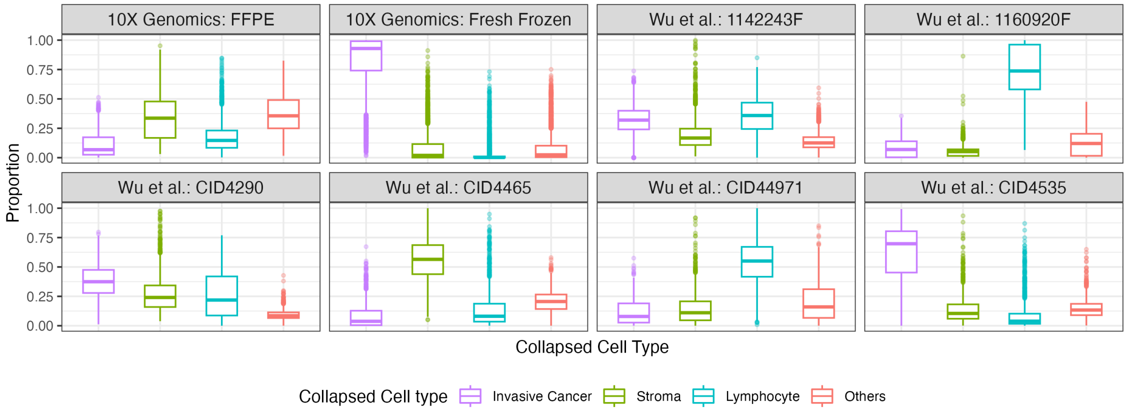

(B) Four collapsed cell types.

Supplementary Figure 5: Boxplot showing the distribution of cell type proportions in all the patches sourced from 10x Genomics and Wu et al. [1], stratified by sample.

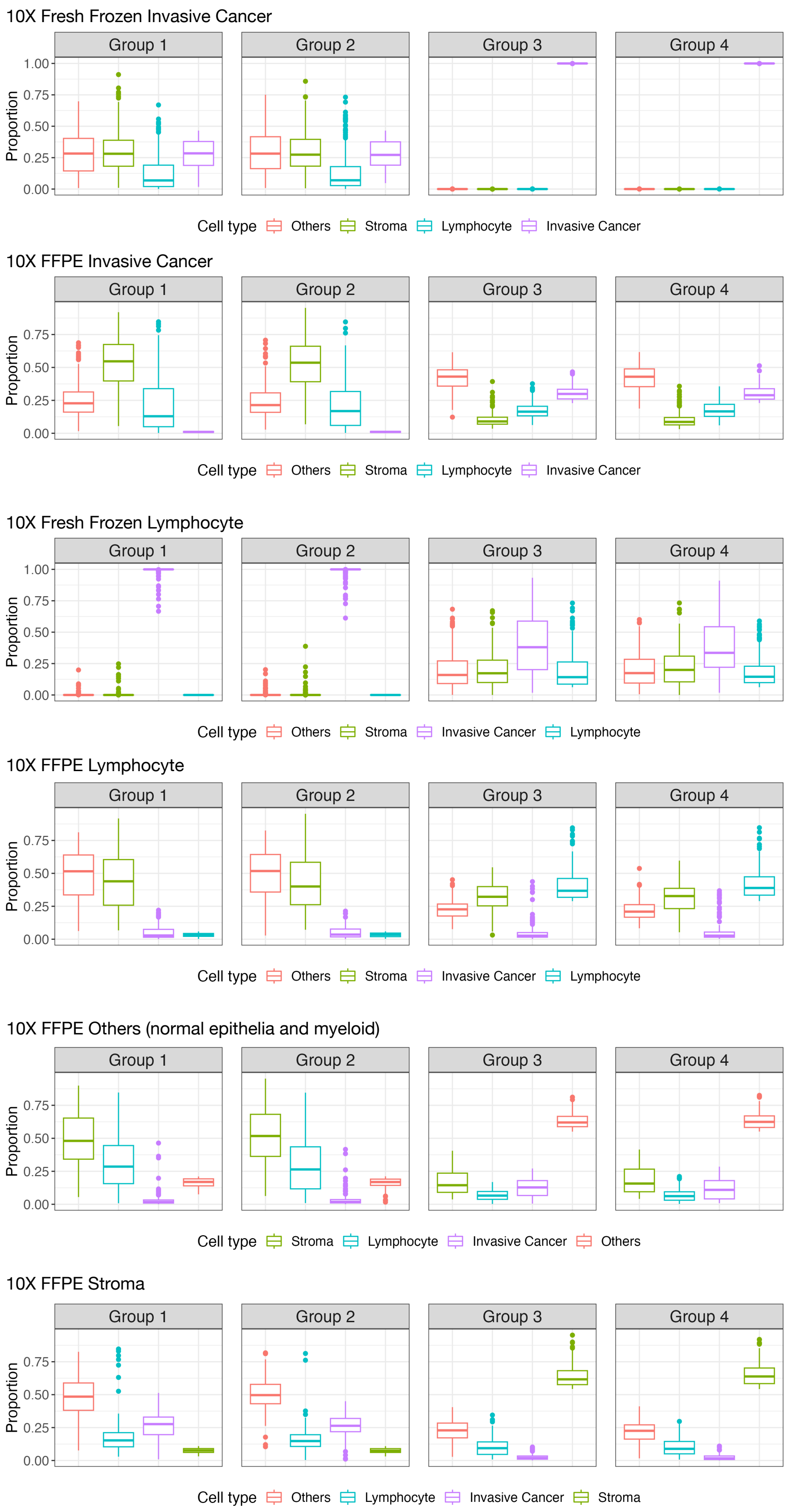

Supplementary Figure 6: Four groups of samples selected to have higher proportions of one cell type in groups 3 and 4, and lower proportions of this cell type in groups 1 and 2, using the data from two 10x samples.

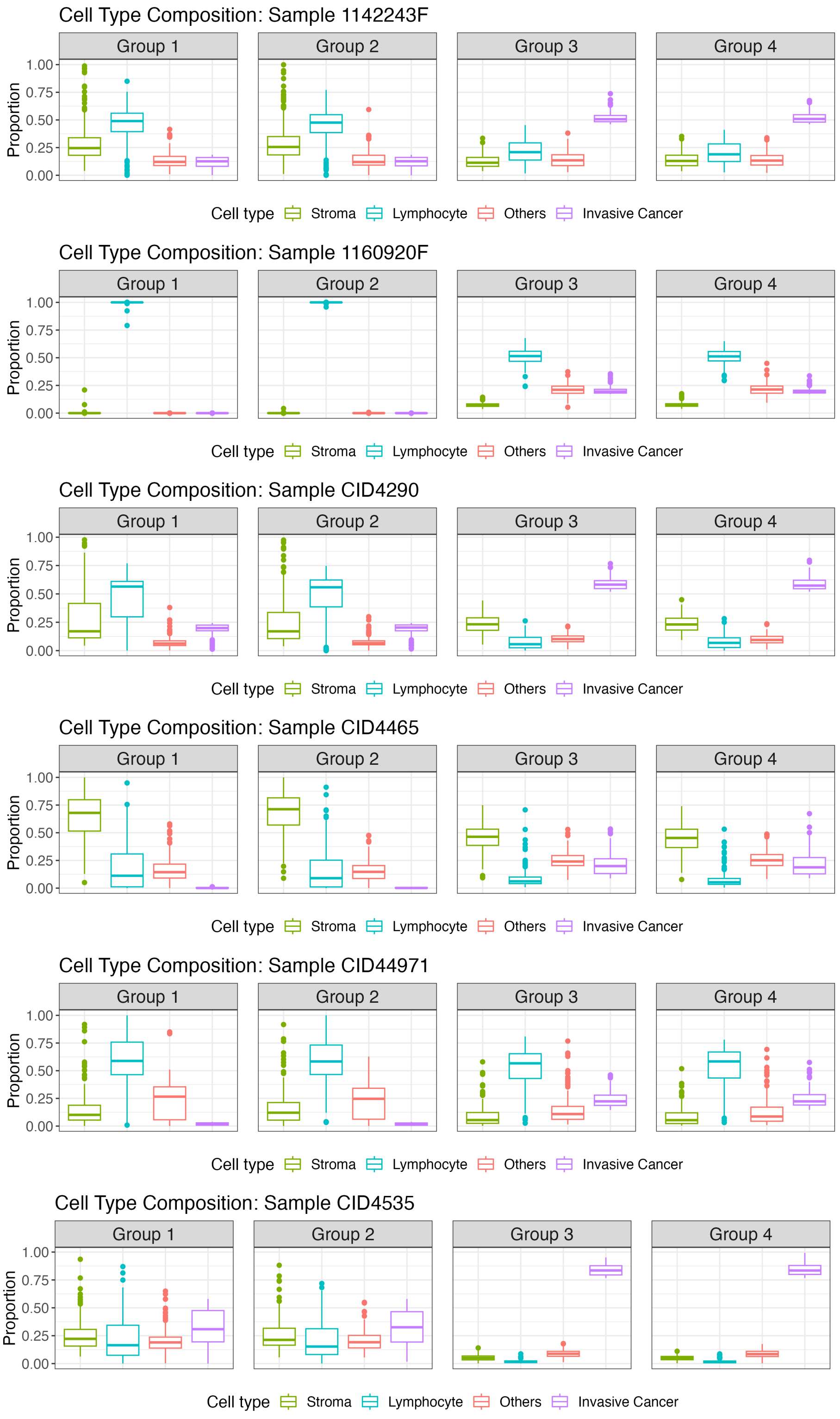

Supplementary Figure 7: Four groups of samples selected to have higher proportions of **invasive cancer** in groups 3 and 4, and lower proportions of invasive cancer in groups 1 and 2, using the 6 samples from Wu et al. [1].

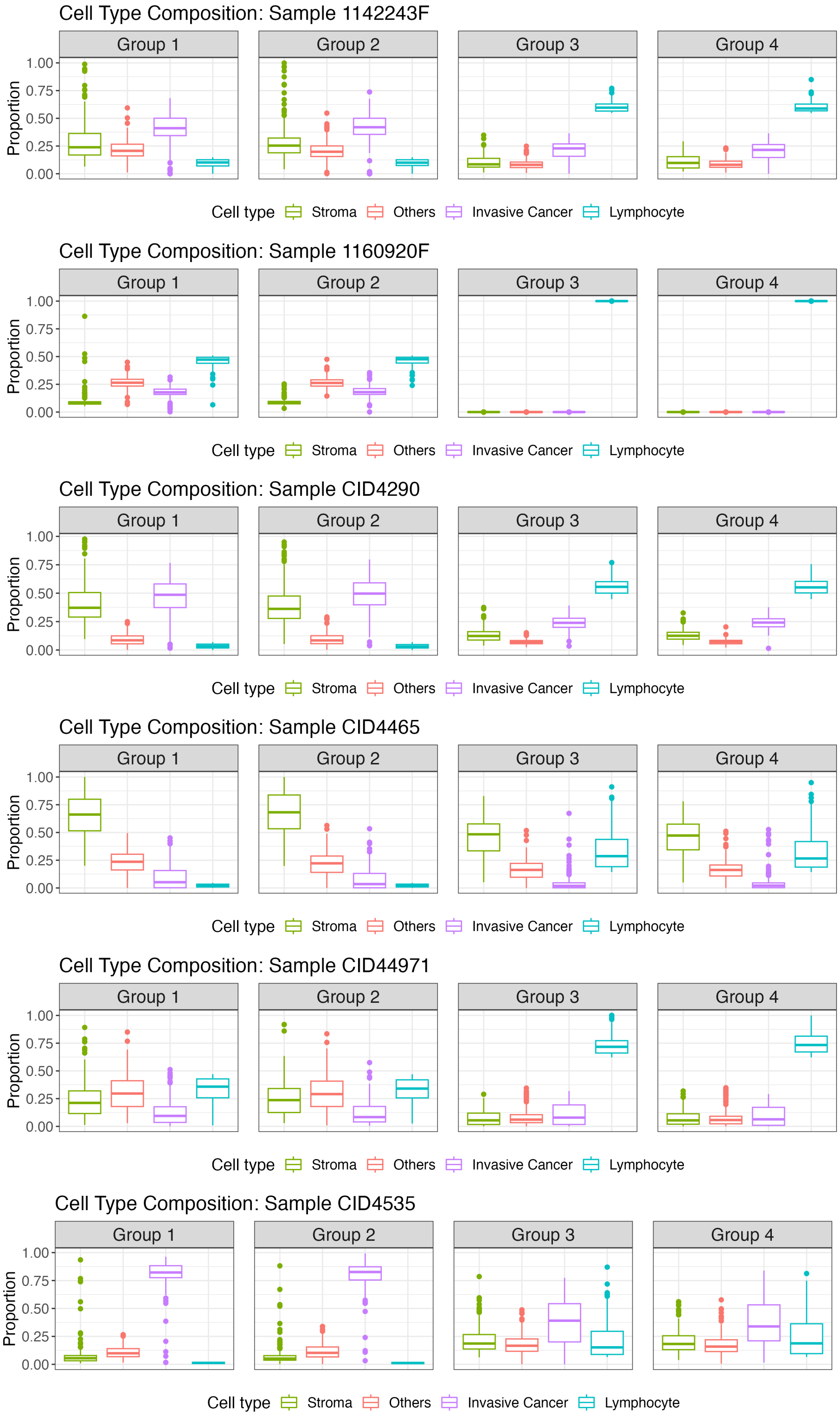

Supplementary Figure 8: Four groups of samples selected to have higher proportions of **lymphocyte** in groups 3 and 4, and lower proportions of invasive cancer in groups 1 and 2, using the 6 samples from Wu et al. [1].

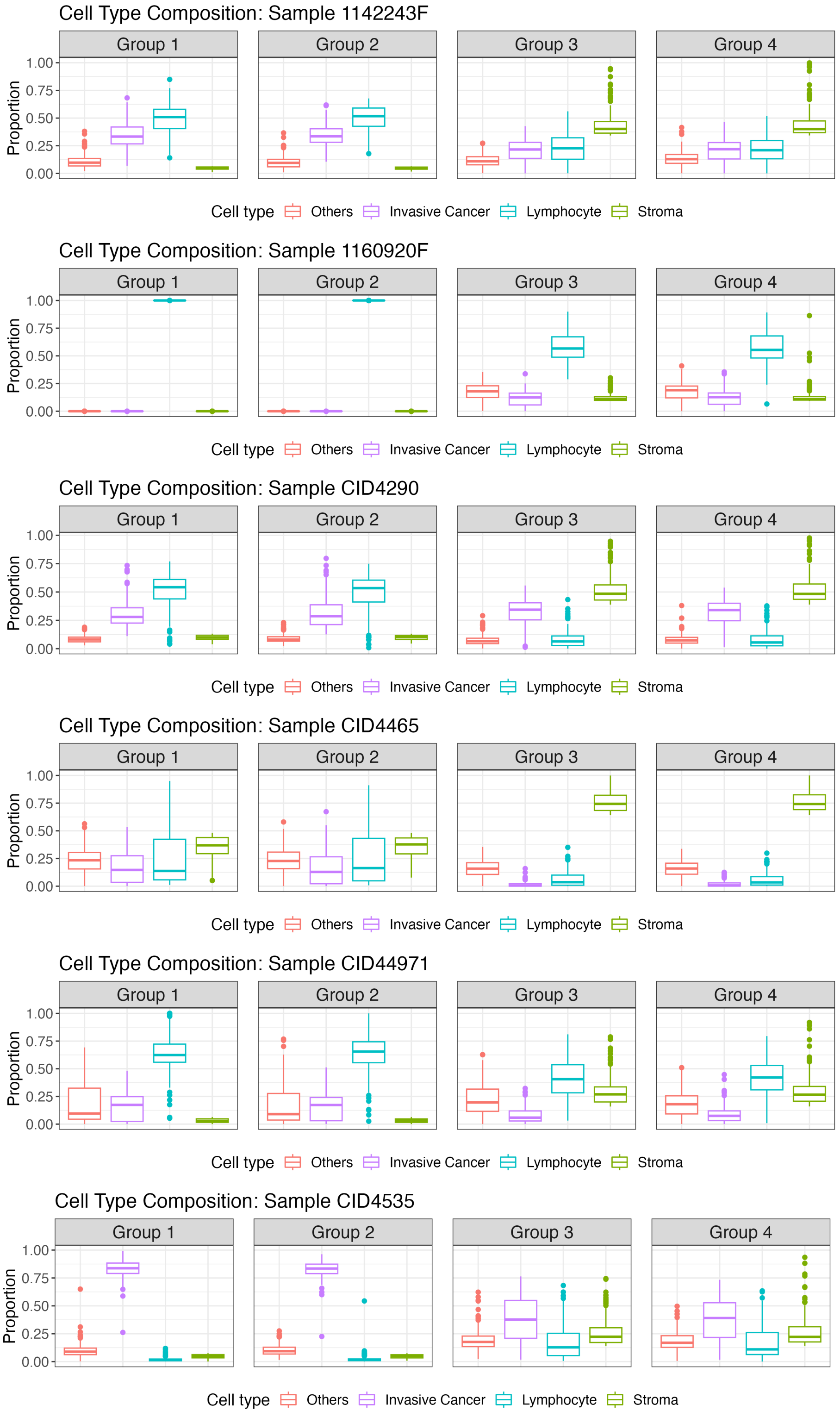

Supplementary Figure 9: Four groups of samples selected to have higher proportions of **stroma** in groups 3 and 4, and lower proportions of invasive cancer in groups 1 and 2, using the 6 samples from Wu et al. [1].

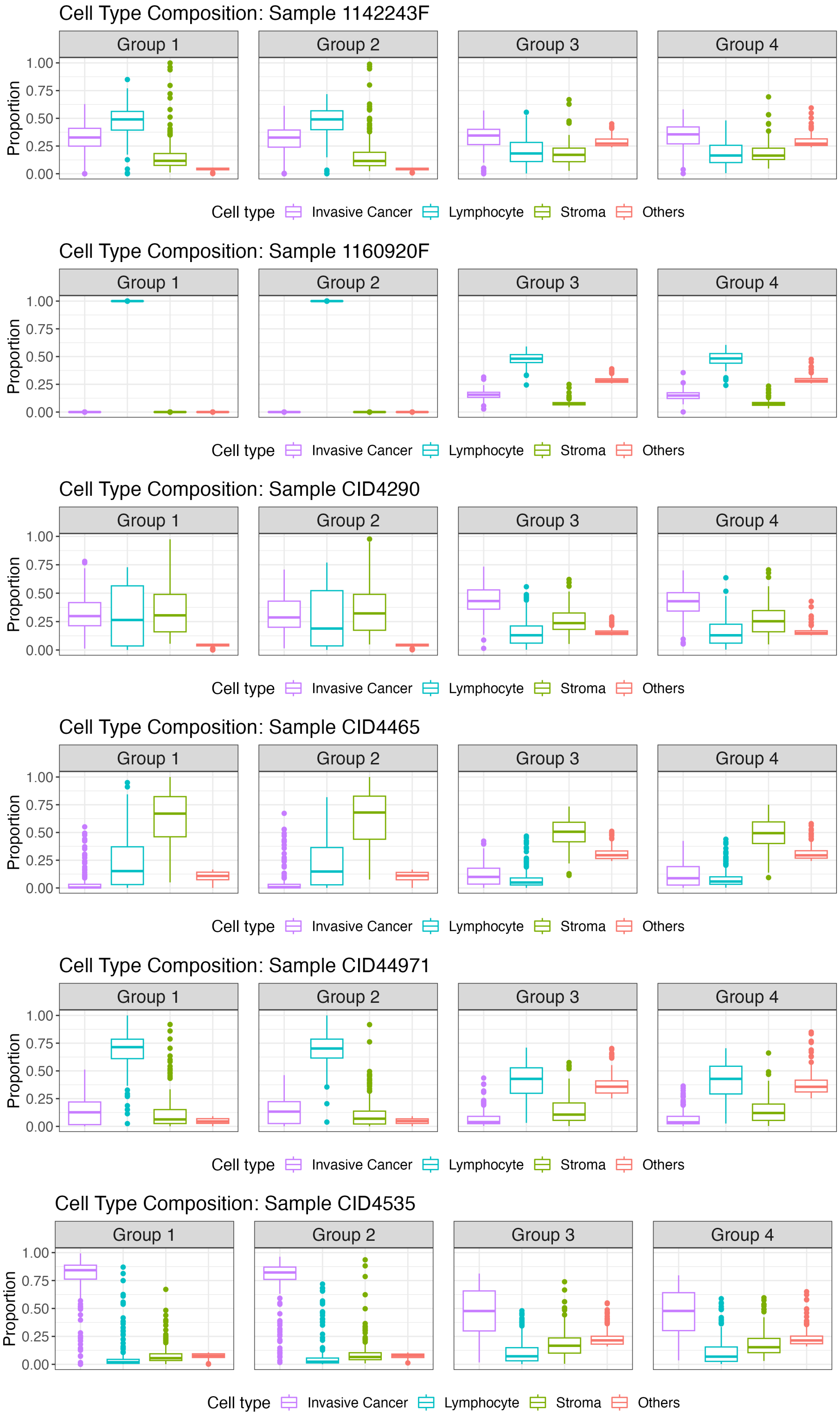

Supplementary Figure 10: Four groups of samples selected to have higher proportions of **other cell types (normal epithelia and myeloid)** in groups 3 and 4, and lower proportions of invasive cancer in groups 1 and 2, using the 6 samples from Wu et al. [1].

#### 2.2 Prediction of cell type proportions using H&E image data

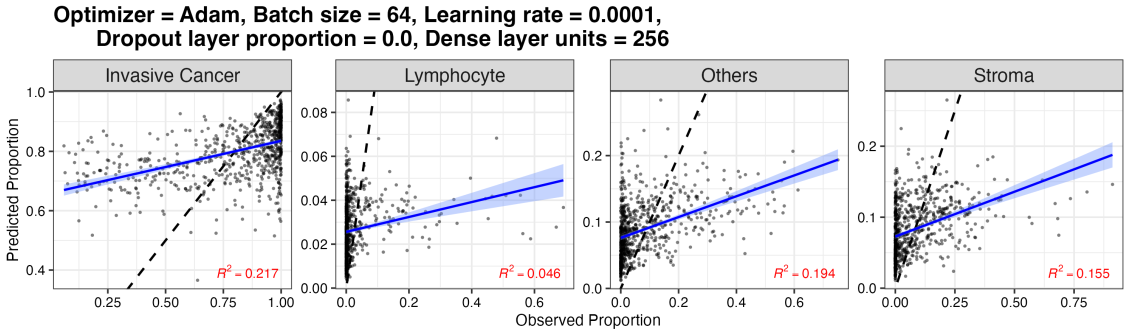

(A) 10x fresh frozen sample.

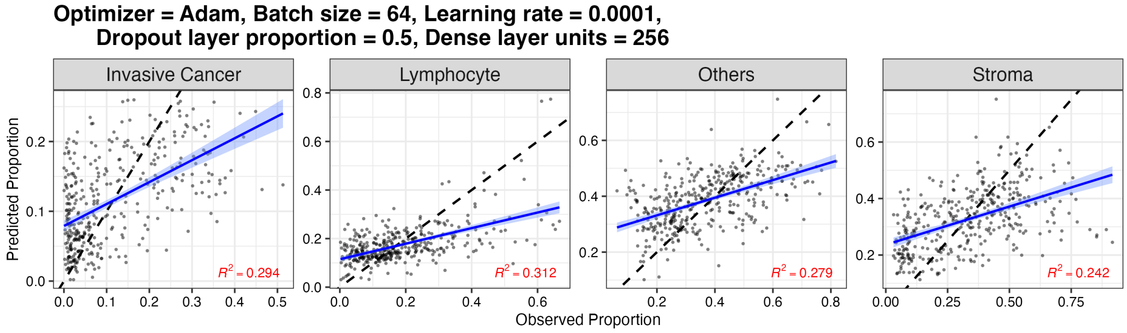

(B) 10x FFPE sample.

Supplementary Figure 11: Predicted cell type proportion in testing data, using neural network trained using the data from the 10x fresh frozen sample and the 10x FFPE sample.

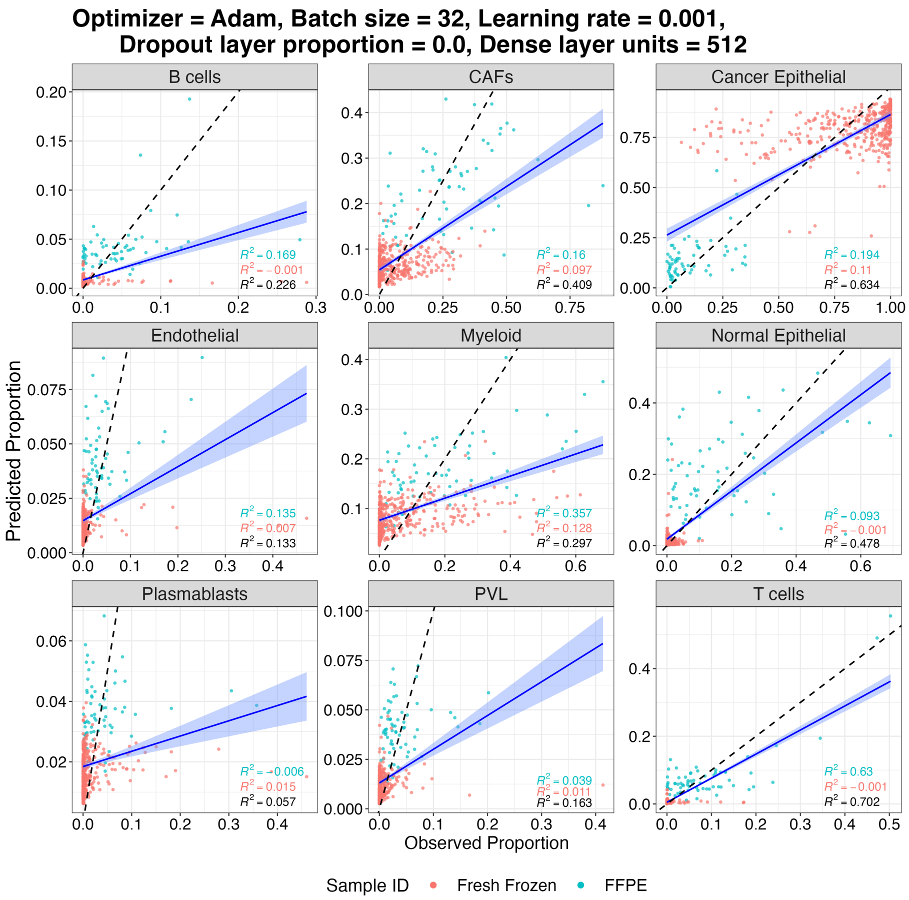

Supplementary Figure 12: Prediction cell type proportions for nine cell types in the testing data of two 10x samples. The title of the plot shows the optimal hyperparameters selected by the validation data.

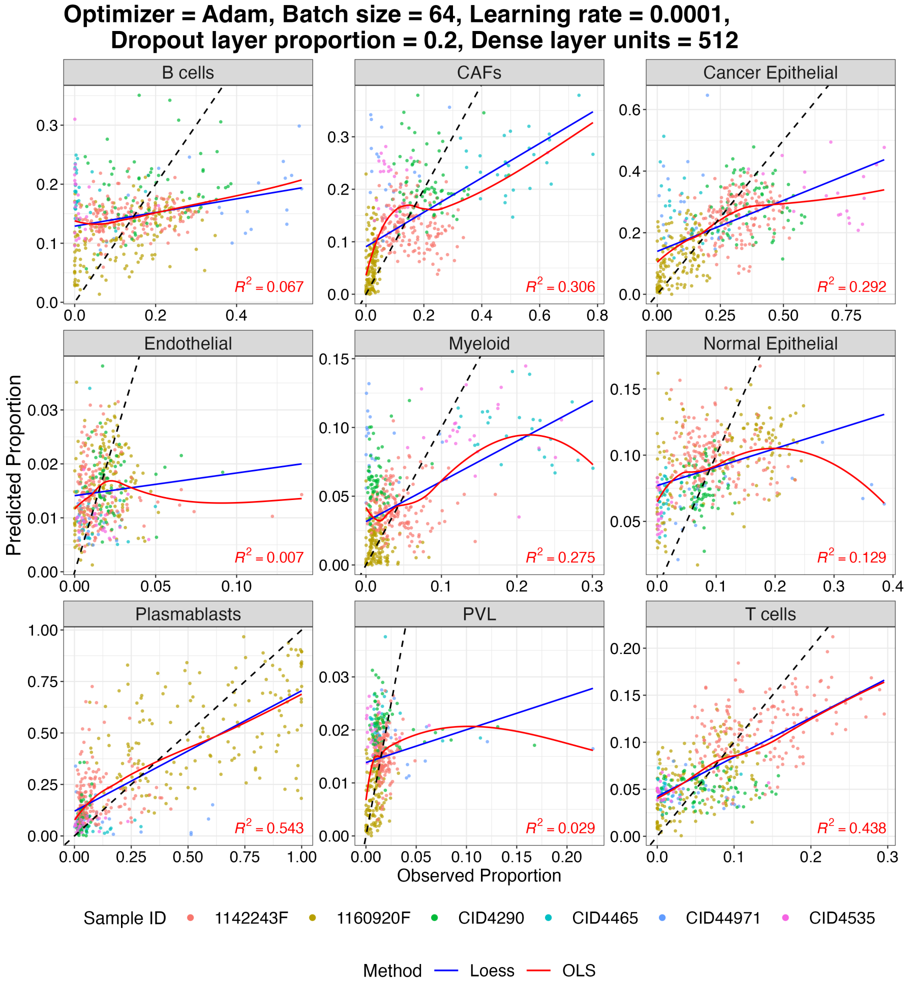

Supplementary Figure 13: Prediction cell type proportions for nine cell types in the testing data of six samples from Wu et al. [1]. The title of the plot shows the optimal hyperparameters selected by the validation data.

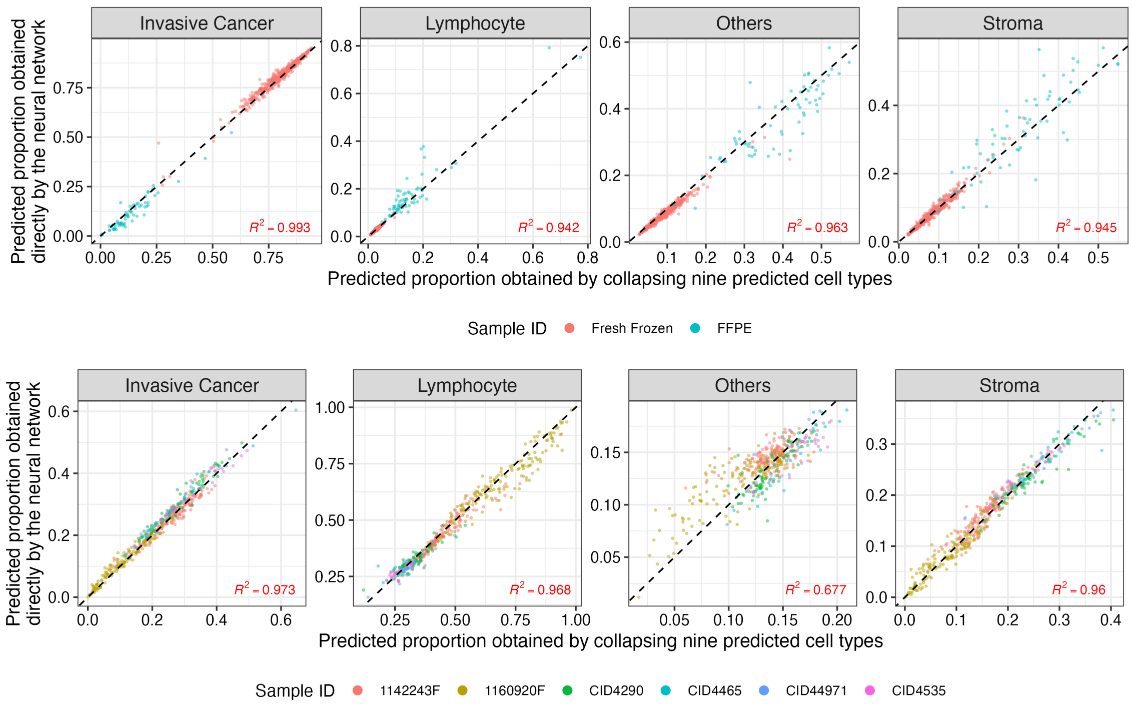

Supplementary Figure 14: Comparisons of the cell type proportions of 4 cell types by two methods: direction predicting the proportions of these four cell types versus predicting 9 cell types and then collapse them to 4 cell types. Upper panel: results for two 10X samples. Lower panel: results for 6 samples from Wu et al. [1].

##### 2.3 Classification using H&E image data

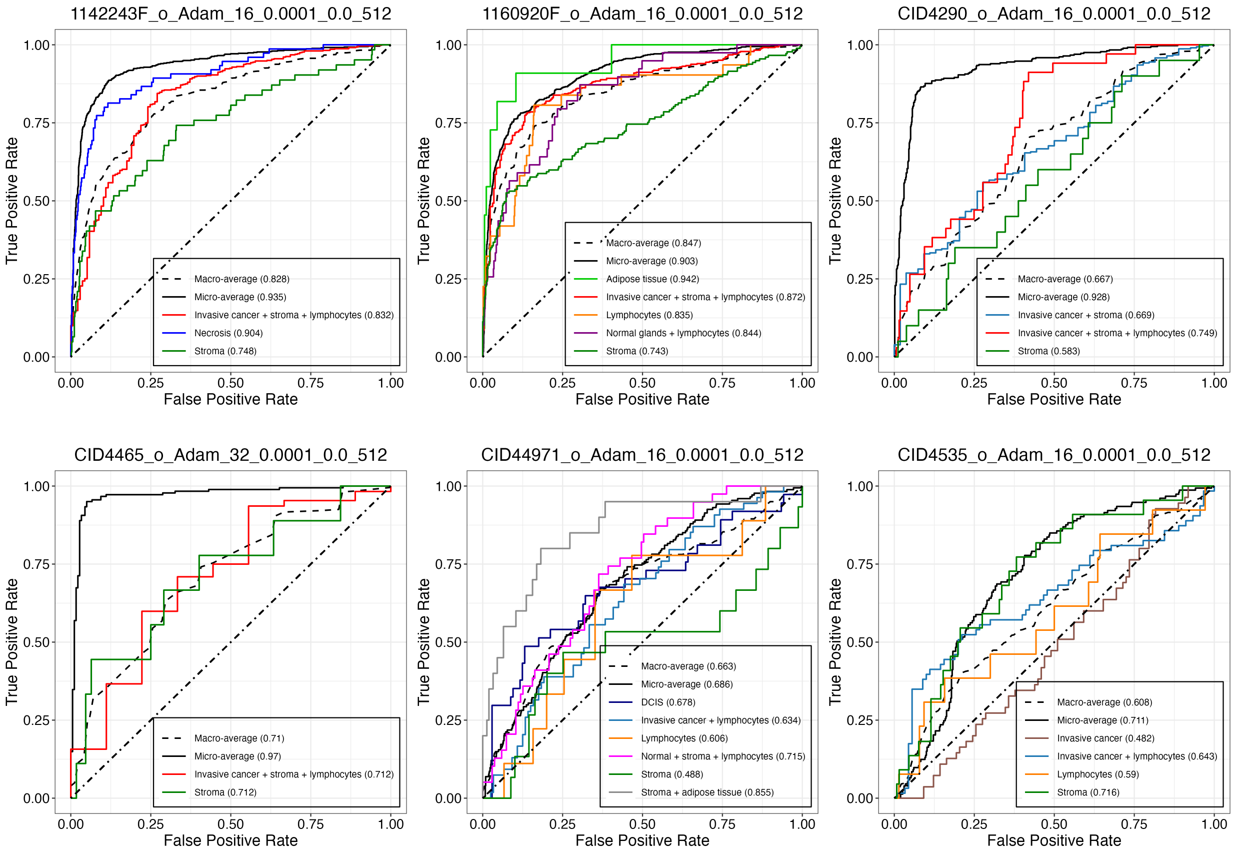

Supplementary Figure 15: ROC curves for the classification task in the testing data of six samples from Wu et al. [1]. The optimal neural network configuration is selected using validation data.

[4] Yuhan Hao, Tim Stuart, Madeline H Kowalski, Saket Choudhary, Paul Hoffman, Austin Hartman, Avi Srivastava, Gesmira Molla, Shaista Madad, Carlos Fernandez-Granda, and Rahul Satija. Dictionary learning for integrative, multimodal and scalable single-cell analysis. *Nature Biotechnology*, 2023.
